# u4atac regulates cilium biogenesis through splicing of the minor intron of *tmem107l* and *rfx7b* in zebrafish developing brain

**DOI:** 10.64898/2026.08.20.745718

**Authors:** Cyril Jovani, Alexia Rabec, Marianne Gaubert, Deepak Khatri, Emma Garnier, Audric Cologne, Anne Meiller, Justine Guguin, Alicia Besson, Sylvie Mazoyer, Marion Delous

**Author notes:** Correspondence to : Marion DELOUS.

## Abstract

Bi-allelic variants of *RNU4ATAC*, transcribed into the minor spliceosome component U4atac snRNA, are associated to variable severity of microcephaly, growth retardation, skeletal dysplasia and immunodeficiency as main features. Previous studies highlighted the dramatic effect of U4atac deficiency on splicing of U12-type introns, which represent less than 1% of all introns in the human genome. More recently, our team evidenced a link between U4atac and the primary cilium/centrosome complex through the identification of patients carrying *RNU4ATAC* bi-allelic variants and exhibiting an atypical Joubert syndrome, a well-known ciliopathy. Yet, the underlying mechanisms remain elusive.

Here, we further explored the link of *RNU4ATAC* to primary cilium and aimed at identifying ciliary U12-type intron containing genes that contribute to the brain abnormalities seen in patients. For that, we performed a transcriptomic analysis of heads of our morpholino oligonucleotide (MO)-mediated u4atac zebrafish model. Through the combined analysis of the generated dataset with those obtained from *RNU4ATAC* patients’ cells, we identified two candidate genes: *TMEM107*, coding for a structural protein of the cilium transition zone, and *RFX7*, encoding a transcription factor involved in primary cilium formation. By conducting complementary genetic approaches in zebrafish model, we showed that both gene orthologues, *tmem107l* and *rfx7b*, functionally interact with u4atac and are required for correct brain development.

Altogether, our findings establish TMEM107 and RFX7 as key components of the molecular pathway linking U4atac dysfunction to ciliary defects and impaired brain development, providing new physiopathological insights and therapeutic perspectives for *RNU4ATAC*-related disorders.

## INTRODUCTION

In eukaryotes, pre-mRNA splicing is carried out by two distinct machineries: the major and the minor spliceosomes, which process U2- and U12-type introns, respectively. The latter are about 850 in the human genome, located in 750 genes, representing less than 1% of all introns (Cologne et al., 2019). Despite their small number, these introns, and the minor spliceosome that excises them, are crucial for development of organisms, as exemplified by various models in which a functional component of the machinery is deficient, whether U11, U12, U4atac or U6atac small nuclear RNA (snRNA) (Baumgartner et al., 2018; Drake et al., 2020; König et al., 2007; Otake et al., 2002; Shikara et al., 2026) or specific proteins such as RNPC3 or ZRSR2 (Markmiller et al., 2014; Weinstein et al., 2022). Notably, in human, deficiency of U4atac snRNA, transcribed from the *RNU4ATAC* gene, causes a subset of rare monogenic disorders, formerly known as Taybi–Linder syndrome (TALS, also known as microcephalic osteodysplastic primordial dwarfism type I, MOPD1) (MIM#210710), Roifman syndrome (RFMN) (MIM#616651), and Lowry–Wood syndrome (LWS) (MIM#226960) (Benoit-Pilven et al., 2020), now collectively referred to as *RNU4ATAC*-opathies (Duker et al., 2023). These disorders share core clinical features, notably primary microcephaly, short stature, skeletal dysplasia and immunodeficiency, even though recent analyses of large cohorts showed that *RNU4ATAC*-opathies consist of a continuum of phenotypes, ranging from microcephalic primordial dwarfism (MPD) (early mortality in >70% of the cases) to pauci-symptomatic cases (Cuinat et al., 2026; Matalon et al., 2026).

In order to understand the impact of U4atac deficiency on U12-type (or minor) intron splicing, several transcriptomics analyses have been performed, mostly in patients’ cells. Altogether, these studies showed massive minor intron retention (IR), i.e. IR in most expressed minor intron-containing genes (hereafter, MIG), at various levels depending on the genes or cell types (Cologne et al., 2019; Dinur Schejter et al., 2017; Merico et al., 2015). Downstream cellular processes that could be altered by these IR have been rarely studied, so that physiopathological mechanisms leading to clinical features seen in *RNU4ATAC*-opathies remain elusive. Our team recently showed that U4atac loss of function results in alteration of cilium function in patients’ fibroblasts as well as in morpholino oligonucleotide (MO)-mediated u4atac zebrafish model, thus explaining the ciliopathy-related phenotypes seen in *RNU4ATAC*-related patients, especially those with atypical Joubert syndrome (JBTS) (Khatri et al., 2023). Cilia are microtubule-based organelles that arise from the mother centriole and are present on the surface of most vertebrate cell types. As key regulators of processes such as cell cycle control, developmental signalling pathway integration (MAPK, Wnt, Hedgehog) and fluid circulation, cilium dysfunction can affect multiple organs and systems, thus leading to a large group of diseases called ciliopathies (Reiter & Leroux, 2017).

Since we showed that 12% of MIG are cilium-related genes, we hypothesised, to explain the link between U4atac deficiency and ciliary defects, that these latter are likely secondary to minor IR (Khatri et al., 2023). To go further on this work and identify ciliary MIG whose expression alteration participates to the phenotypes seen in patients, we conducted a transcriptomic analysis in u4atac-deficient zebrafish embryos. By comparing the results with previous published RNAseq datasets from patients’ fibroblasts (Cologne et al., 2019), we identified two ciliary candidate MIG associated with neurodevelopmental disorders, *TMEM107* and *RFX7* (Chinen et al., 2022; Harris et al., 2021; Lambacher et al., 2015; Ledger et al., 2023; Manojlovic et al., 2014), that exhibit significant minor IR both in zebrafish and humans. Using epistasis and rescue approaches in zebrafish embryos, we showed that these genes are involved in the phenotypes of u4atac-deficient models, particularly microcephaly. Hence, our findings suggest that *TMEM107* and *RFX7* may largely contribute to the pathophysiological mechanisms leading to microcephaly in *RNUATAC* patients.

## Material and methods

### Cell culture

Fibroblasts were cultivated in HAMF10 medium (Eurobio, CM1H1000-01), complemented with 12% fetal bovine serum (FBS, Eurobio, CVFSVF00-01) and 1% penicillin-streptomycin (PS, Gibco, 15140122). Human telomerase-immortalized retinal pigment epithelial cells (hTERT-RPE1) were grown in Dulbecco’s modified Eagle DMEM/F12 medium (Thermofisher, 31330095), supplemented with 10% FBS and 0.01mg/mL hygromycin B. Lymphoblastoid cell lines (LCL) were cultivated in RPMI medium (MP Biomedicals, 0910601-CF) with 10% FBS. Induced Pluripotent Stem Cells (iPSC) issued from a healthy European male (PCi-CAU2) (Phenocell, Grasse, France) were cultivated in 35-mm vitronectin-coated dish (STEMCELL Technologies, 7180) with mTeSR Plus medium (STEMCELL Technologies, 100–0276) supplemented with 10 μM ROCK inhibitor Y-27632 (STEMCELL Technologies, 72302) for thawing, and then with 0,1% PS. Differentiation of iPSC into neuronal stem cells (NSC) and neurons was conducted as previously described (Guguin et al., 2024).

### Zebrafish husbandry and micro-injections of embryos

Zebrafish (*Danio rerio*) adults and embryos were maintained at 28.5°C under standard conditions (Kimmel et al., 1995). The following strains were used: mixed wild-type ABxTU and *Tg(wt1b:EGFP)li1* (Perner et al., 2007). Occasionally, 1-phenyl-2-thiourea (PTU) 0.003% (w:v) was added in E3 medium at 24 hpf to inhibit pigmentation, thereby facilitating pronephros visualization. Organs from one-year-old adult fish (equal ratio of males and females) were dissected following the procedure described in Gupta & Mullins, 2010.

Morpholino oligonucleotides (MO) targeting *rfx7b*, *tmem107l, tmem216*, *cep290* were designed and provided by Gene Tools, LLC (Philomath, USA) (Supp Table 1). u4atac MO and its five-mismatch control MO were described previously (Khatri et al., 2023). MO were injected into the yolk at one-cell stage (3 to 10 ng per embryo, 0.4 ng for *u4atac* MO for loss-of-function experiments; 0.3 to 0.7 ng per embryo, 0.1 ng for u4atac MO for epistasis experiments).

CRISPR RNA (crRNA) sequences were designed using the online tool CRISPOR (https://crispor.gi.ucsc.edu/) (Supp Table 1). crRNA and tracrRNA (IDT, 1077024) were diluted to 3 µM in IDTE buffer, denaturated at 95°C for 5 min, and cooled down to room temperature to allow guide RNA (gRNA) duplex formation. CRISPR/Cas9 RNPs were then formed with combination of gRNA with 0.5 µg Cas9 protein (IDT, 1081060) at a 2:3 ratio (v/v), incubated at 37 °C for 10 min, and stored at –20°C overnight prior to injections. 1 nL of this mix was injected into each embryo.

For rescue experiments, *TMEM107* mRNA was synthesised from *TMEM107* Human tagged clone (Origene, RC223559), using mMESSAGE mMACHINE T7 kit (Invitrogen, AM1344), followed by RNA purification with RNA clean and concentrator kit (Zymo Research, R1017). 180 pg were co-injected with u4atac MO or *tmem107l* MO per embryo.

For RNAseq experiment, human U4atac snRNA molecules were synthesised using the MAXIscript T7 Transcription Kit (ThermoFisher Scientific) as previously described (Khatri et al., 2023), and injected into the yolk at one-cell stage (0,25 ng MO + 65 pg snRNA).

### DNA and RNA extraction, RT-PCR and RT-qPCR

Genomic DNA (gDNA) from zebrafish embryos was extracted by treating samples with Proteinase K (1 mg/mL) in TE buffer at 65°C for 4 hours followed by an inactivation at 90°C for 10min.

For RNAseq, total RNA was extracted from three independent pools of 20 heads sectioned from zebrafish embryos at 48 hours post-fertilization (hpf) using TRI reagent (Zymo Research, R2050-1-200) and following manufacturer’s instructions. RNA was then purified using RNA clean and concentrator kit (Zymo Research) prior to be treated with DNase I (Invitrogen, 3644103). RNA integrity number was > 9 for all samples. For RT-PCR and RT-qPCR, total RNA from zebrafish embryos (pool of 20), organs (pool of two adults) and human cells was extracted and treated with DNase I using the “NucleoSpin RNA Plus” kit (Macherey-Nagel) or “RNeasy Mini” kit (Qiagen, 74104).

For cDNA synthesis, 1.5 μg of total RNA was reverse-transcribed using GoScript Reverse Transcriptase kit (Promega, A5003) with 50 ng of random hexamers or oligo(dT) primers. PCR was performed using GoTaq Green Master Mix (Promega, M7122), and quantitative PCR with OneGreen Fast qPCR premix (Ozyme, OZYA008-200XL) using Rotor-Gene Q (Qiagen) following manufacturers’ instructions. RT-qPCR reactions were performed in triplicates with the appropriate primers (0.5 µM) (Supp. Table 1), using *RPS17* as the reference gene for human cells and *gapdh* for zebrafish embryos. Relative expression levels were calculated using the 2^-ΔΔCt method. Intron retention (IR) was quantified as the ratio of unspliced transcript level to the total of unspliced plus spliced transcript form levels, after normalization to *RPS17/gapgh* gene level.

### cDNA library preparation and high-throughput RNA sequencing

Four hundred nanograms of RNA from 48 hpf heads of MO-injected embryos (with or without human U4atac snRNA) were sent for RNA-sequencing to IntegraGen Genomics (Evry, France), where a DNA library was generated with the “TruSeq Stranded mRNA Sample Prep” kit (Illumina) that comprises a step of mRNA purification using oligo(dT) beads. RNA-seq experiments have been performed on a HiSeq 4000 sequencer (Illumina), yielding approximately 2049 million of stranded two time 100 bp paired-end reads.

### Bioinformatic analyses

For identification of minor introns in the zebrafish genome, MIG were identified using two tools: Tyler Alioto’s scoring matrices and script, originally developed for the generation of the U12db (Alioto, 2007), and intronIC v1.4.0 (Moyer et al., 2020). For intronIC, the —allow_multiple_isoforms parameter was enabled to classify introns across all annotated isoforms per gene. Although intronIC directly classifies introns as major or minor using a support vector machine (SVM) algorithm, it also computes, for each intron, the probability of being a minor intron. Following the recommendations of the authors, introns with a probability ≥ 90% were considered minor introns. Both tools were used on the Ensembl 110 GRCz11 genome annotation. The final list of minor introns is the union of the results obtained by the three methods: Tyler Alioto, intronIC SVM and intronIC minor intron probability (Supp. Dataset 1). Gene orthology of zebrafish genes with human was determined using biomaRt v2.62.1 on the Ensembl 110 GRCz11 and GRCh38 genome annotations (Durinck et al., 2005, 2009).

For the analysis of MIG expression throughout zebrafish development, we used the publicly available RNAseq dataset of White et al. (2017) (ENA, ERP014517). It consists of a total of 90 samples, with five biological replicates for each of 18 developmental stages (1-cell, 2-cell, 128-cell, 1k-cell, dome, 50% epiboly, shield, 75% epiboly, 1–4 somites, 14–19 somites, 20–25 somites, prim-5, prim-15, prim-25, long pec, protruding mouth, day 4, and day 5 post-fertilization), sequenced on an Illumina HiSeq 2500 platform using 100 bp paired-end reads (White et al., 2017). FastQC v0.12.1 (available at https://www.bioinformatics.babraham.ac.uk/projects/fastqc/) and Cutadapt v4.9 (Martin, 2011) were used to perform quality control and adapters trimming, followed by alignment of reads on the GRCz11 zebrafish reference genome using STAR v2.7.11a (Dobin et al., 2013). Expression levels of genes were quantified using RSEM v1.3.3 (Li & Dewey, 2011). Genes with estimated length < 1 bp were removed. Next, for each developmental stage, genes were considered expressed with a mean TPM ≥ 1 across the five biological replicates. Only protein coding genes were analysed. Gene expression trajectories throughout development were analysed using ImpulseDE2 (Fischer et al., 2018). Default parameters were used, except for the boolIdentifyTransients option, which was set to TRUE to enable the identification of transiently activated or deactivated genes. Significant genes were defined with a False Discovery Rate (FDR) < 0.01.

For analysis of RNAseq dataset issued from u4atac MO-injected samples, pre-processing of data was done as mentioned above, and genes were considered expressed with a mean TPM ≥ 1 in at least one triplicate. Differential gene expression analysis was performed using DESeq v1.42.1 (Love et al., 2014) with a FDR < 0.05 and genes with an absolute fold change (FC) greater than 2 (|log2(FC)| ≥ 1) were considered significantly differentially expressed (DE). Intron retention analysis was performed using IRFinder-S v2.0.1 (Lorenzi et al., 2021). Only introns located within the previously defined expressed genes were included in the analysis. Introns were not considered if the LowCover tag was present in at least 2 samples in a triplicate, and we kept only introns with 0.05 ≤ IR ratio ≤ 1 in at least one triplicate. Differential intron retention analysis was performed using the IRFinder diff mode, which employs the DESeq2 algorithm with the cooksCutoff and independentFiltering options enabled. Introns with a FDR < 0.05 and an absolute delta IR ratio (mean IR ratio condition – mean IR ratio control) |ΔIR ratio| ≥ 0.1 were considered significantly differentially spliced (DS).

For principal component analysis (PCA), we used the prcomp package function from the R package v1.7-11 (https://github.com/sdray/ade4) (Bougeard & Dray, 2018) on VST for expression level and intron retention analysis. The most variable values (up to 500) were used (as conventionally done in DESeq2) and the first (PC1) and second (PC2) most explanatory axes were plotted.

Gene ontology (GO) Enrichment Analysis was used for functional annotation and pathway analysis with TopGO v2.58.0 (Alexa et al., 2006) R tool, with the default “weight01” algorithm, the Fischer test and the org.Dr.eg.db annotation database for zebrafish genes. Genes without GO annotation were excluded. Only pathways with p-value < 0.01 were considered.

Euler plots were generated with eulerr v7.1.0 (Larsson & Gustafsson, 2018), and graphical representations of bioinformatics datas were produced with ggplot2 v3.5.2 (Wickham, 2016).

### Phenotypic analysis

For phenotype assessment, zebrafish embryos at 48 hpf were anesthetized with MS222 0.02% (Sigma Aldrich) and photographed using Leica M165 FC stereomicroscope. Images were analyzed using ImageJ software. Body curvature was quantified by the ventral bending angle and categorized as severe (<90°), moderate (90–120°), mild (120–160°), or absent (160–180°). Microcephaly and microphthalmia were assessed using the head area-to-body length and eye area-to-body length ratios, respectively.

### Whole-mount in situ hybridization and O-dianisidine staining

Whole-mount *in situ* hybridization was carried out on embryos from 1 to 4 dpf following standard protocols (Thisse & Thisse, 2008; primers in Supp. Table 1).

O-dianisidine staining, used to detect hemoglobin in embryos, was performed according to previously described method (Detrich et al., 1995). Embryos were dechorionated at 48 hpf and stained for 15 minutes in the dark in solution containing O-dianisidine 0.6 mg/mL, 0.01 M sodium acetate (pH 4.5), 0.65% H_2_O_2_, and 40% ethanol. Then, embryos were fixed in 4% PFA for 20 minutes, dehydrated in methanol then bathed in a clearing solution of benzyl benzoate/ benzyl alcohol (2:1, vol/vol) for imaging.

### Statistical analysis

All the data are reported as the mean with standard deviation (SD) or median with 95% confidence interval (CI) of at least three independent experiments. Normality of datasets was evaluated using the Shapiro-Wilk test, and outliers identified with the Grubb’s test. All hypothesis tests were two-sided, and statistically significant differences (p-value < 0.05) were calculated by one-way ANOVA or t-tests as indicated in figure legends. If datasets did not follow normal distribution or if the sample size was too small, a non-parametric test (Kruskal-Wallis or Mann-Whitney) was used. Statistical analyses were performed using GraphPad Prism software.

## Results

### Characterization of zebrafish minor intron-containing genes and of their expression throughout development

The zebrafish displays a high degree of conservation of the minor spliceosome (Baumgartner et al., 2019; Moyer et al., 2020; Weinstein et al., 2022; Olthof et al., 2022). Using two different algorithms, U12db (Alioto, 2007) and IntronIC (Moyer et al., 2020), we identified by summing up the results 743 U12-type introns located in 677 genes annotated in the GRCz11 genome assembly (615 genes containing a single U12-type intron, 58 containing two and 4 containing three). Among them, 565 genes (83.4%) have at least one human orthologue that contains a U12-type intron, 34 have human orthologue(s) without U12-type intron, while 78 (11.5%) have no human orthologue (Supp. Dataset 1). As in humans, gene ontology (GO) term analysis based on the 634 MIGs *vs* the 17,083 U2 genes (genes with only U2-type introns) with GO annotation highlighted biological processes associated to membrane depolarization and ion homeostasis among the most significant (Figure 1A, Supp. Dataset 1). Of note, GO terms related to primary cilium formation and function (“positive regulation of smoothened signaling pathway”) are also enriched in zebrafish MIGs.

**Figure 1.**
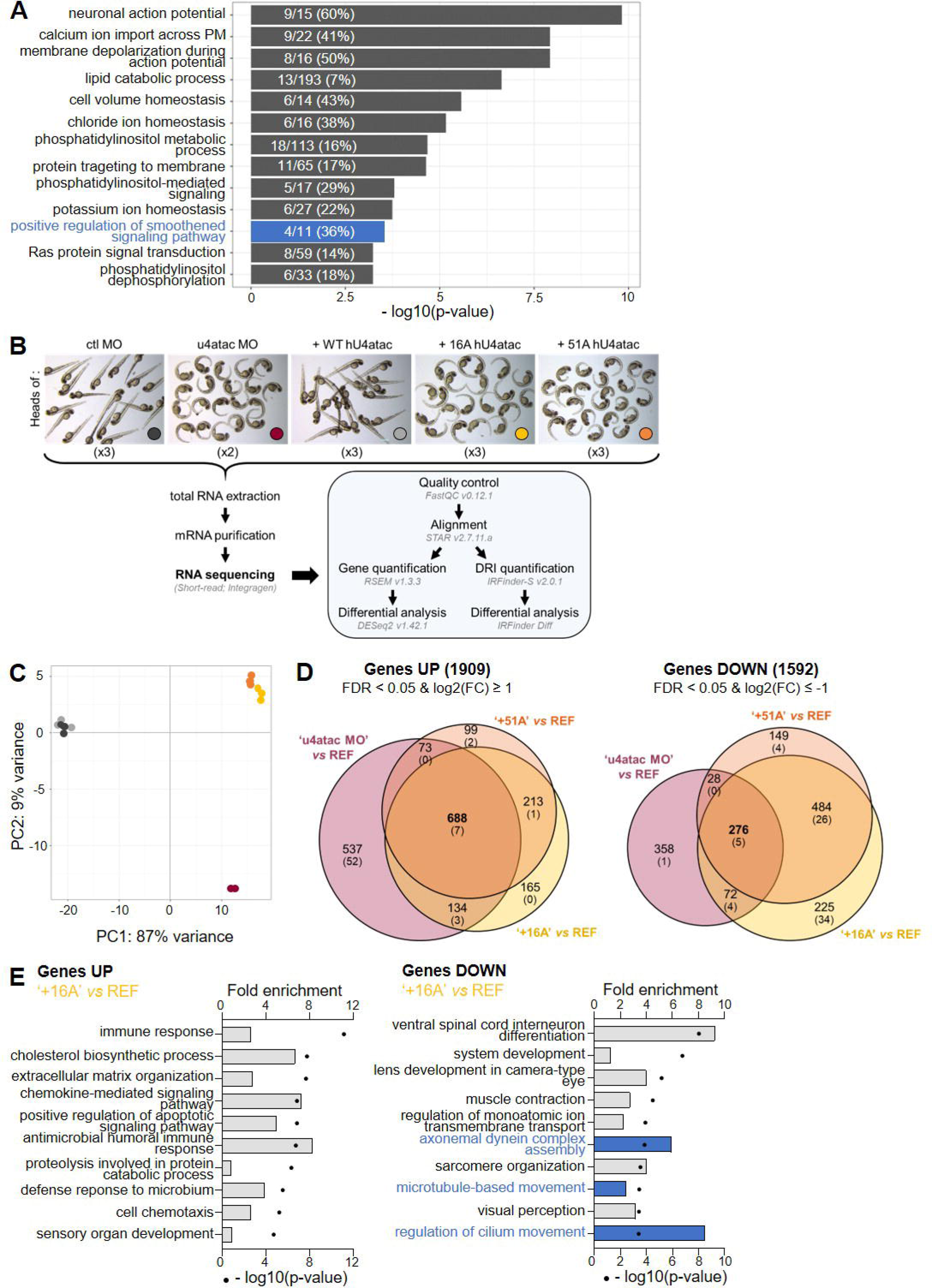
Co-injection in zebrafish of u4atac MO and U4atac-16A or -51A snRNA expression results in transcript level alteration of genes associated to immune response and cilium biogenesis. **(A)** GO term enrichment analysis of the 634 MIGs compared with the 17,083 U2 genes annotated with GO terms in GRCz11 zebrafish genome assembly. Numbers indicate the count of enriched genes relative to the total number of genes associated with the GO term. Cilium-related term is highlighted in blue. **(B)** Workflow of the RNAseq analysis performed on zebrafish heads collected at 48 hpf. For each condition, three biological replicates consisting of a pool of 20 zebrafish heads were analyzed, except for the u4atac MO condition (two replicates). **(C)** Principal component analysis (PCA) of gene expression profiles across the five experimental conditions, based on the 500 most variable genes. Same color legend as in B. **(D)** Euler diagrams showing the numbers of up- and down-regulated genes relative to the REF condition (FDR < 0.05 and |log2(FC)| ≥ 1) across the different experimental conditions. In brackets, the number of MIGs only. **(E)** GO term enrichment analysis of up- and down-regulated genes in +16A condition compared to the reference, ranked according to the adjusted p-value (black dots). Bars represent fold enrichment for each GO term. PM, plasma membrane; DSI, differentially spliced intron; PR, photoreceptor.

The majority of these MIGs (64%) are expressed across all embryonic and larval stages from 1-cell stage to 5 dpf (mean TPM ≥ 1 in at least one developmental stage), based on the expression dataset of White et al., 2017. Globally, they cluster into four different expression patterns: they are either down-(13%), up-(21%), or transiently down-(31%) or up-(35%) regulated during the time course (Supp. Fig. 1). Altogether, these observations indicate that MIGs are widely expressed during development in zebrafish.

### u4atac deficiency in zebrafish developing brain leads to U12-type intron retention with small impact on transcript levels

To identify the genes contributing the most to *RNU4ATAC*-related brain anomalies, we performed a bulk short-read RNA sequencing on 3 independent pools of 20 zebrafish embryo heads at 48 hpf. Five conditions were compared: control MO (ctl), u4atac MO, and rescue with either human wild-type (+WT), n.16G>A (+16A) or n.51G>A (+51A) U4atac snRNA (Figure 1B), the phenotypes of which were previously described in Khatri et al., 2023.

Principal component analysis (PCA) based on the expression of the 500 most variable genes showed that u4atac MO, +51A and +16A conditions were highly distinct from ctl MO and +WT conditions according to PC1 (87% of variance), with much less difference between u4atac MO and +51A or +16A conditions (PC2 at 9%) (Figure 1C). It is noteworthy that control MO and +WT conditions looked undistinguishable, which was further demonstrated by the absence of differentially expressed or spliced genes (Supp. Fig. 2A-B), as well as by the full rescue of morphological phenotype in +WT condition (Khatri et al., 2023; Figure 1B). The two conditions were thus pooled into a single reference group (hereafter, REF) for the subsequent analyses. Overall, we identified about 2,150 (10 to 12%) differentially expressed (DE) genes with FDR < 0.05 and |log2(FC)| ≥ 1 between each condition group (u4atac MO, +16A, +51A) and the reference, with a higher proportion of up-regulated genes in u4atac MO condition (Figure 1D, Supp. Dataset 2). Pooling all u4atac-deficient conditions together, it represents a total of 3,501 (19%) DE genes. Considering MIGs only, DE genes represented from 12% (u4atac MO) to 7% (+51A) of the expressed MIG genes, depending on the condition, so the expression of this category of genes was not particularly impacted by u4atac deficiency (Supp. Dataset 2). As shown by the PCA, the two conditions expressing the variants (+16A and +51A) were much more similar, with very few differences between them, than the u4atac MO condition (Figure 1D, Supp. Figure 2C, Supp. Dataset 2). GO term analyses of DE genes revealed an enrichment for immune response processes among up-regulated genes for each of the three conditions (u4atac MO, +16A, or +51A *vs* REF) (Supp. Fig. 1D, Figure 1E). Among down-regulated genes, the enriched terms are neuronal development in u4atac MO condition (Supp Fig. 1D, Supp. Dataset 2), and muscle formation and cilium function in +16A or +51A conditions (Figure 1E, Supp. Dataset 2).

We then analysed IR across all experimental conditions using IRFinder-S tool, which allowed to analyse 533 U12-type introns. The PCA, based on the 500 most variable U12-type introns, showed a clear separation of u4atac deficient conditions u4atac MO from the reference (83% of variance), with a less clear separation for +16A, and especially for +51A (Figure 2A). Indeed, u4atac MO condition exhibited a much higher proportion of U12-type introns with significant IR (FDR < 0.05) – 96 % *vs* 73% (+16A) or 58% (+51A) – and among those, a higher proportion of U12-type introns with ΔIR ratio > 0.1 (92% *vs* 58% (+16A) or 46% (+51A)) (Figure 2B, Supp. Dataset 3). In addition, merely all genes with U12-type retained introns in +16A and +51A conditions are common to u4atac MO condition (Figure 2C). Hence, injection of u4atac MO leads to more global and stronger U12-type intron retentions than 16A or 51A variants, indicating that these variants are hypomorphic. It is noteworthy that 88.7% of MIGs with U12-type IR are not differentially expressed (Supp. Dataset 4), suggesting a moderate impact of IR on the level of expression, as we previously observed in patients’ cells (Cologne et al., 2019).

**Figure 2.**
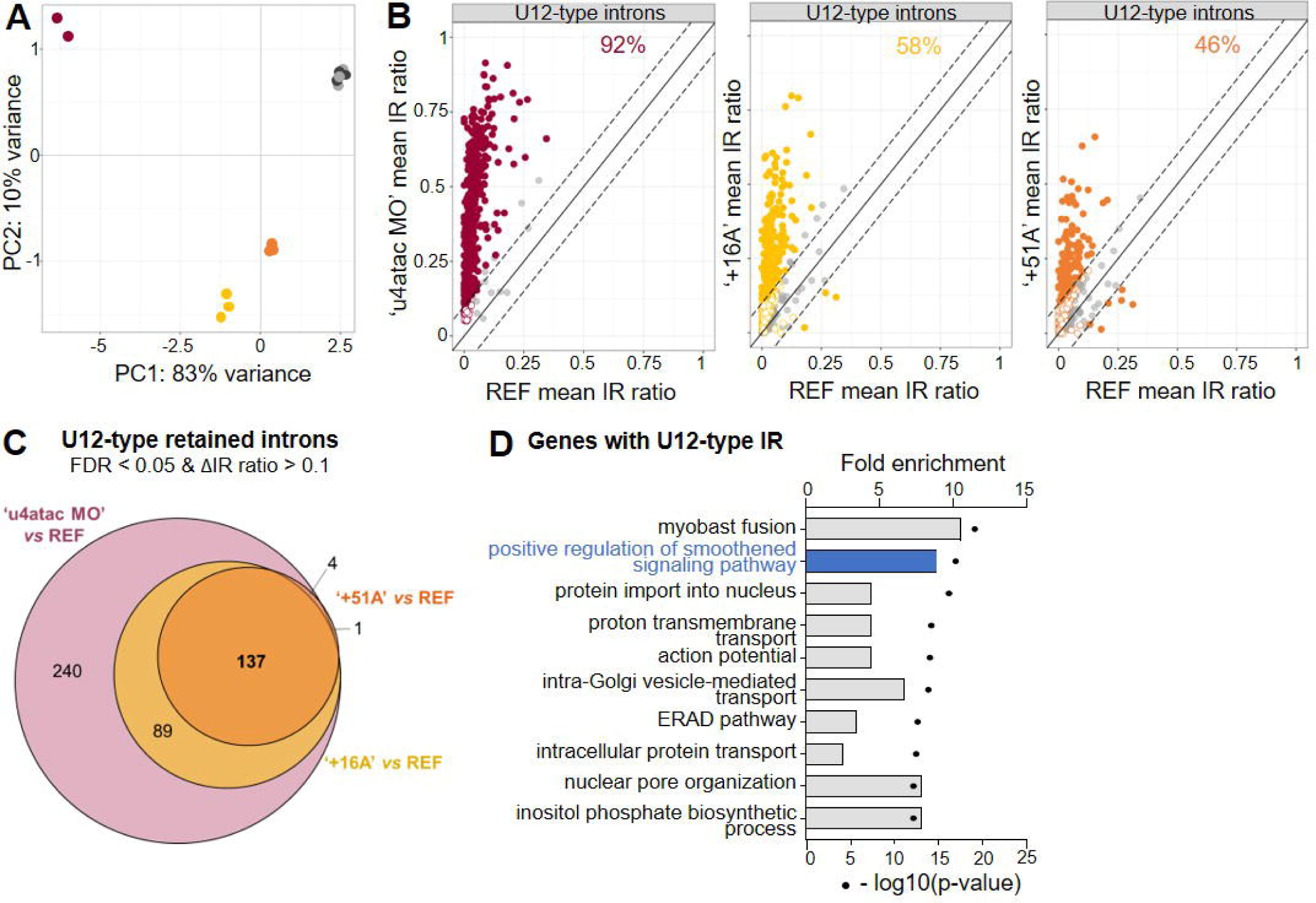
u4atac deficiency leads to massive U12-type intron retention in cilium-related MIGs. **(A)** PCA of U12-type intron retention levels across the five experimental conditions, based on the 500 most variable U12-type introns. Same color legend as in Figure 1B. **(B)** Distribution of U12-type intron retention levels in each condition relative to the REF condition. All 533 detected U12-type introns are shown. The solid diagonal line indicates equal retention levels between the two conditions (y = x). Dashed lines delineate a ±10% difference threshold relative to the equality line. Colored dots indicate statistically significant differences (FDR < 0.05), with plain dots being introns with ΔIR ratio > 0.1 and circles with ΔIR ratio < 0.1, while grey dots correspond to introns with non-significant differences (FDR > 0.05). Percentages indicate numbers of U12-type introns with significant ΔIR ratio > 0.1 over all significant U12-type introns. **(C)** Euler diagram showing significant (FDR < 0.05) U12-type retained introns with a ΔIR ratio > 0.1 relative to the reference (REF). **(D)** GO term enrichment analysis of genes containing U12-type intron retention in u4atac deficient conditions, seen in C, ranked according to the adjusted p-value (black dots). Bars represent fold enrichment for each GO term. ERAD, endoplasmic reticulum-associated protein degradation.

Curiously, when analysing U2-type IR, we observed that both +16A and +51A conditions displayed a higher splicing efficiency of about 16% of detected introns (FDR < 0.05 and ΔIR ratio < -0.1); the same trend is also seen in u4atac MO condition, but it concerns only 1.6% of detected introns (Supp. Fig. 3, Supp. Dataset 3). We interpreted this effect as a decrease of physiological U2-type IR, a known form of alternative splicing involved in the fine regulation of transcript levels (Boutz et al., 2015; Braunschweig et al., 2014; Grabski et al., 2021). It is noteworthy that the large majority (> 96%) of these better spliced U2-type introns are within genes with U2-type introns only.

To identify the cellular processes that are possibly the most altered by the U12-type intron retentions, we performed a GO term enrichment analysis on all MIGs (440) with significant and differentially retained U12-type introns in u4atac deficient conditions compared to the reference. Among the enriched cellular processes, the second most enriched pathway is associated to primary cilium function, while several others are linked to intracellular protein transport (Figure 2D, Supp. Dataset 3). Considering the 86 cilium-related MIG we identified (Khatri et al., 2023), ciliary genes overall represent 14.5% of all 440 genes with significant U12-type IR (ΔIR ratio > 0.1).

#### TMEM107 *and* RFX7 *as candidate MIGs to explain ciliary phenotypes*

To select relevant ciliary MIGs to explore further, we crossed our dataset of zebrafish genes with U12-type IR with that issued from TALS or JBTS patients’ cells we previously published (Cologne et al., 2019; Khatri et al., 2023) (Supp. Dataset 5). Among the 361 zebrafish genes with transcripts with U12-type IR and at least one orthologue in human, 219 (60.7%) are found impacted in human cells (FDR < 0.05, ΔIR ratio > 0.1), 28 of them being cilium-related genes (Figure 3A, Supp. Dataset 5). Among them we focused on genes associated with a pathology (n=12), and further on those with ubiquitous or brain expression, which led us to select *TMEM107* and *RFX7*. *TMEM107* encodes a transmembrane protein of the cilium transition zone associated to several ciliopathies, i.e. Joubert, Meckel and Oro-Facial-Digital syndromes (Lambacher et al., 2015; Shaheen et al., 2015; Shylo et al., 2016). *RFX7* codes for a transcription factor of the RFX family, known to regulate ciliary gene expression during development (Chu et al., 2010). In particular, for RFX7, it indirectly regulates ciliogenesis in *X. laevis* neural tube by controlling *rfx4* expression (Manojlovic et al., 2014). D*e novo* variants in *RFX7* have been identified in patients with neurodevelopmental disorders, with or without microcephaly (Harris et al., 2021; Ledger et al., 2023; Lee et al., 2021; Sisroe et al., 2024).

**Figure 3.**
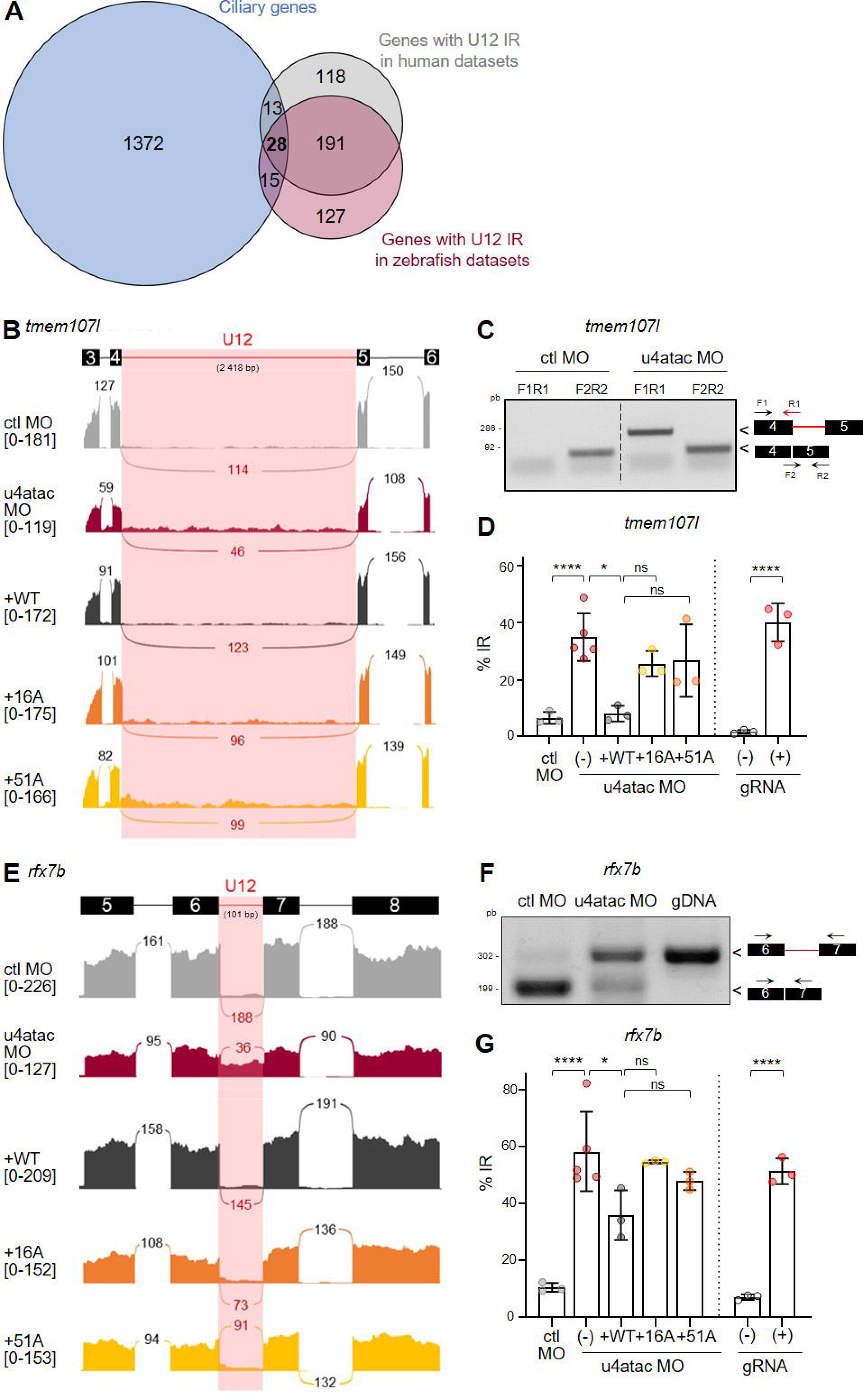
*tmem107l* and *rfx7b* U12-type introns are mis-spliced in u4atac-deficient zebrafish embryos. **(A)** Euler diagram illustrating the overlap of transcripts with U12-type IR between ciliary MIGs (Khatri et al., 2023) and genes with U12-type IR in zebrafish dataset (present study) and human dataset (Cologne et al., 2019). **(B, E)** Sashimi plots of U12-type intron splicing (highlighted in pink) of *tmem107l* (B) and *rfx7b* (E) across the different experimental conditions. On the lines, the number of reads split across splice junctions; in brackets, the range of coverage of each base of the depicted region. **(C, F)** RT-PCR analysis of U12-type intron retention in *tmem107l* (C) and *rfx7b* (F) in control and u4atac MO conditions. Genomic DNA (gDNA) is used as control in F. Primer positions are indicated on the right panel. **(D, G)** RT-qPCR analyses of *tmem107l* (D) and *rfx7b* (G) U12-type intron retention showing the mean ± SD of at least three independent experiments in morphants (u4atac MO) and crispants (gRNA). ns, non-significant, *P<0.05, ****P<0.0001 using one-way ANOVA test with Dunnett’s multiple comparisons (MO conditions), or unpaired t-test (gRNA conditions).

*TMEM107* and *RFX7* have both two orthologues in zebrafish: *tmem107* and *tmem107-like* (*tmem107l*), and *rfx7a* and *rfx7b*, with *tmem107l* and *rfx7b* being their closest human orthologues based on amino acid identity of their functional domains (Supp. Fig. 4A, B). As a first step, we validated the U12-type IR of both *tmem107l* and *rfx7b*, and of their paralogues, in u4atac deficient zebrafish embryos by visualising RNAseq data with IGV and by performing RT-PCR and RT-qPCR (Figure 3B-G, Supp. Fig. 4C-F). U12-type IR has also been confirmed in patients’ fibroblasts for both human orthologues by RT-qPCR (Khatri et al., 2023; Supp. Fig. 4G). As a second step, we validated that *tmem107l* and *rfx7b* are expressed in brain during zebrafish development, using public datasets (DanioCell, Farrell et al., 2018; Sur et al., 2023), as well as *in situ* hybridisation and RT-qPCR approaches (Supp. Fig. 5A-D). In human cells, we showed by RT-qPCR that *TMEM107* is expressed at similar levels in all tested cell types (blood, skin fibroblasts, retinal and brain cells), while *RFX7* is more expressed in these latter (retinal RPE1 cells, iPSC-derived neural progenitors and neurons) (Supp. Fig. 5E).

Taken together, these data identified *TMEM107* and *RFX7* as strong candidate MIGs that may contribute to the physiopathological mechanisms leading to brain anomalies seen in *RNU4ATAC*-associated diseases.

#### Knockdown of tmem107l and rfx7b induces pleiotropic phenotypes in zebrafish, including microcephaly

To demonstrate the involvement of *TMEM107* and *RFX7* in *RNU4ATAC*-related physiopathology, we first assessed the impact of loss of function of *tmem107l* and *rfx7b* on zebrafish development. For doing so, we used MO targeting the exon/U12-type intron junction (hereafter, U12 MO) to attempt to elicit U12-type IR and mimic the loss of function of u4atac, which proved successful (Supp. Fig. 6A-B, D-E). We confirmed the specificity of the MO by using a second MO (Supp. Fig. 6A-B, D) targeting either another splice site (sp MO) for *tmem107l*, or the translation site (ATG MO) for *rfx7b*, as it is maternally contributed (Supp. Fig. 5A) and thus fully spliced in the early stages. Additionally, we used CRISPR/Cas9 RNP injections (gRNA) to induce indels in the *tmem107l* and *rfx7b* coding sequences (Supp. Fig. 6A, C, D, F) and study F0 crispant phenotypes. We previously reported that zebrafish u4atac morphants show ciliopathy-related phenotypes, including ventral body curvature, otolith defects and pronephric cysts, as well as brain haemorrhages and microcephaly (Khatri et al., 2023). We further confirmed these phenotypes using CRISPR/Cas9 RNP targeting *rnu4atac* (Supp. Fig. 7A, B), and noticed that otolith defects were less frequent than in morphants, whereas microcephaly was more penetrant, body curvature more severe, and an additional ciliopathy-related feature, hydrocephaly, was seen (Supp. Fig. 7C-F). Of note, we validated in this *rnu4atac* crispant model by RT-qPCR analysis that *tmem107l* and *rfx7b* U12-type IR was at the same level of that observed in u4atac morphants (Figure 3D-G). As expected given the involvement of *TMEM107* in ciliopathies, we observed ciliary-associated phenotypes in *tmem107l* deficient embryos, including body curvature, pronephric cysts and hydrocephaly (Figure 4A-C). As for *rfx7b* loss of function, hydrocephaly was the only phenotype suggesting ciliary defects, pronephric cysts and otolith defects being occasional, and body curvature mild at most (Figure 4D-F). This result may be explained by the indirect role of rfx7 in ciliary gene regulation, which was demonstrated in *X. laevis* to be dependant of *rfx4* (Manojlovic et al., 2014). In addition to ciliopathy-related phenotypes, we noticed alteration of yolk extension in both *tmem107l* and *rfx7b* deficient animals, as well as reduced number of red blood cells for *rfx7b* (Supp. Fig. 8A-C). Very interestingly, across all experimental conditions, depletion of *tmem107l* or *rfx7b* consistently resulted in marked microcephaly and in microphthalmia (Figure 4A-B, D-E, G, Supp. Fig. 8D), underscoring the critical role of these genes in early brain development in zebrafish.

**Figure 4.**
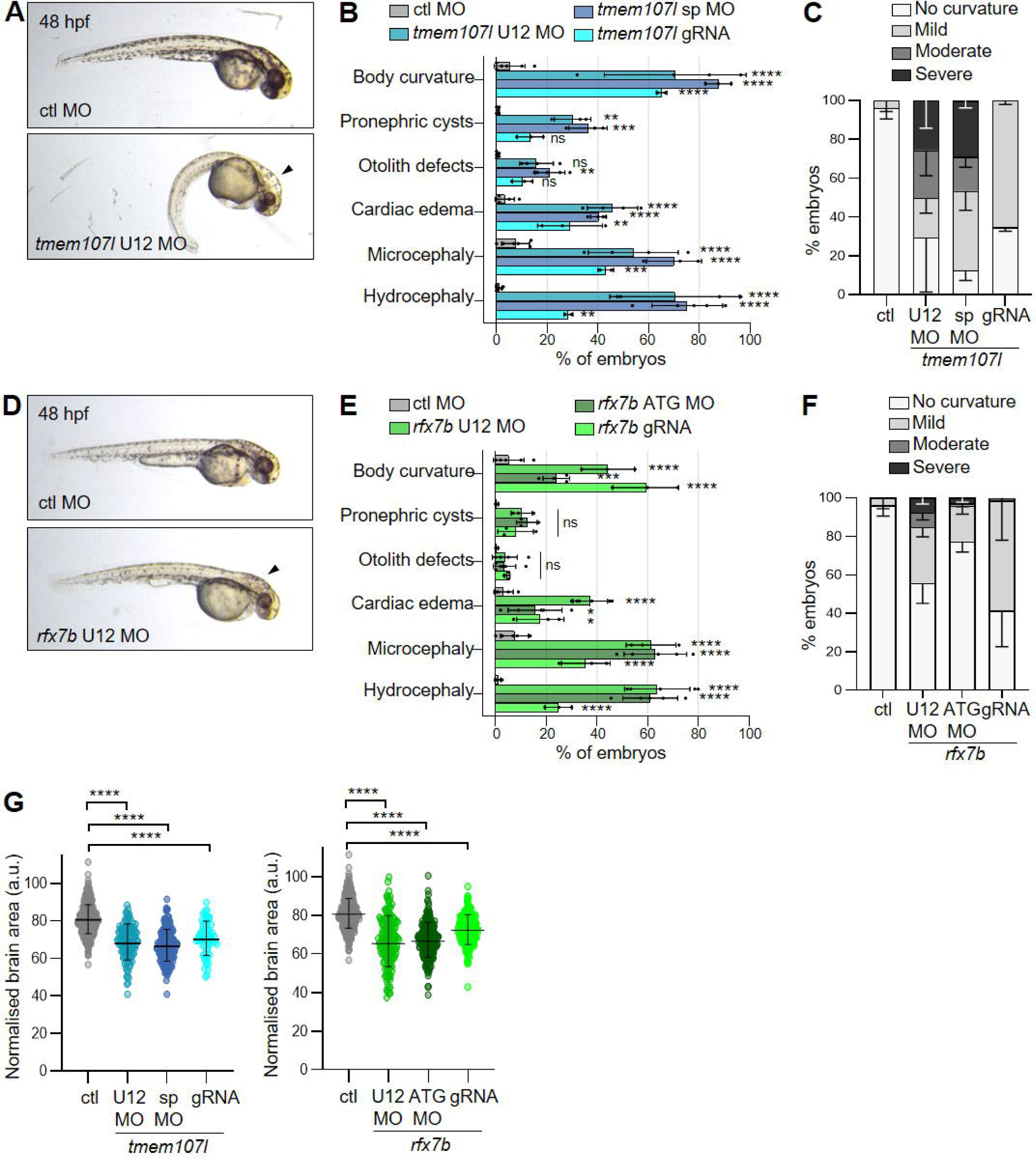
Loss of function of *tmem107l* and *rfx7b* results in ciliopathy-related phenotypes and severe microcephaly. **(A, D)** Representative images of 48 hpf embryos injected with control MO and *tmem107l* (A) or *rfx7b* (D) U12 MO. Black arrows indicate hydrocephaly. **(B, E)** Percentage of embryos displaying each of the observed phenotypes in *tmem107l* (B) or *rfx7b* (E) deficient conditions (using either splice (sp) or ATG MO, U12 MO, or gRNA) compared with control morphants. Graphs represent the mean ± SD of at least three independent experiments, with a minimum of 30 embryos per experiment. ns, non-significant, *P<0.05; **P<0.01; ***P<0.001; ****P<0.0001, following two-way ANOVA with Tukey’s multiple comparison test compared to control MO. N.D., not determined. **(C, F)** Graph showing the distribution of ventral body curvature phenotypes in *tmem107l* (C) and *rfx7b* (F) deficient conditions at 48 hpf, categorized into four classes of severity (no curvature, mild, moderate, and severe) as shown in Supplementary Fig. 7F. Graphs show the mean ± SD from at least three independent experiments, each including a minimum of 30 embryos per batch. **(G)** Graphs showing the normalised head area to total body length in *tmem107l* (left) or *rfx7b* (right) deficient embryos at 48 hpf. Graphs represent the mean ± SD of three independent experiments, with 30 to 60 embryos per experiment. ****P<0.0001, following Kruskal-Wallis test, with Dunn’s comparison test.

It is noteworthy that for both genes, MO-mediated retention of U12-type intron induced phenotypes of comparable severity to those observed in other strategies of knockdown (sp MO, ATG MO or gRNA), indicating that failure to excise U12-type intron likely prevents functional protein synthesis and consequently, correct neurodevelopment.

Altogether, these findings demonstrated that *tmem107l* and *rfx7b* are essential for neurodevelopment and cilium function, and highlighted the critical requirement of the removal of their U12-type intron from pre-mRNA for the genes to fulfil their function. Hence, their subsequent loss of function, downstream U12-type intron retention, may contribute to the phenotypes we observe in u4atac-deficient animals.

#### *u4atac and* tmem107l/rfx7b *are involved in a common process leading to microcephaly*

To further demonstrate the involvement of *TMEM107* and *RFX7* in *RNU4ATAC*-opathy pathogenesis, we performed epistasis analysis using sub-optimal doses (SOD) of *tmem107l* or *rfx7b* U12 MO in combination with SOD of u4atac MO.

First of all, we showed that SOD of each MO induced no detectable phenotype when injected together with the control MO. Conversely, co-injection of u4atac MO SOD with either *tmem107l* or *rfx7b* MO SOD resulted in marked microcephaly, indicating functional interaction between *rnu4atac* and each candidate gene for this phenotype (Figure 5A-B, D-E). We also observed a functional interaction concerning body axis curvature with *tmem107l*, suggesting that this gene and *rnu4atac* are interrelated in cells lining the central canal (Figure 5C, F).

**Figure 5.**
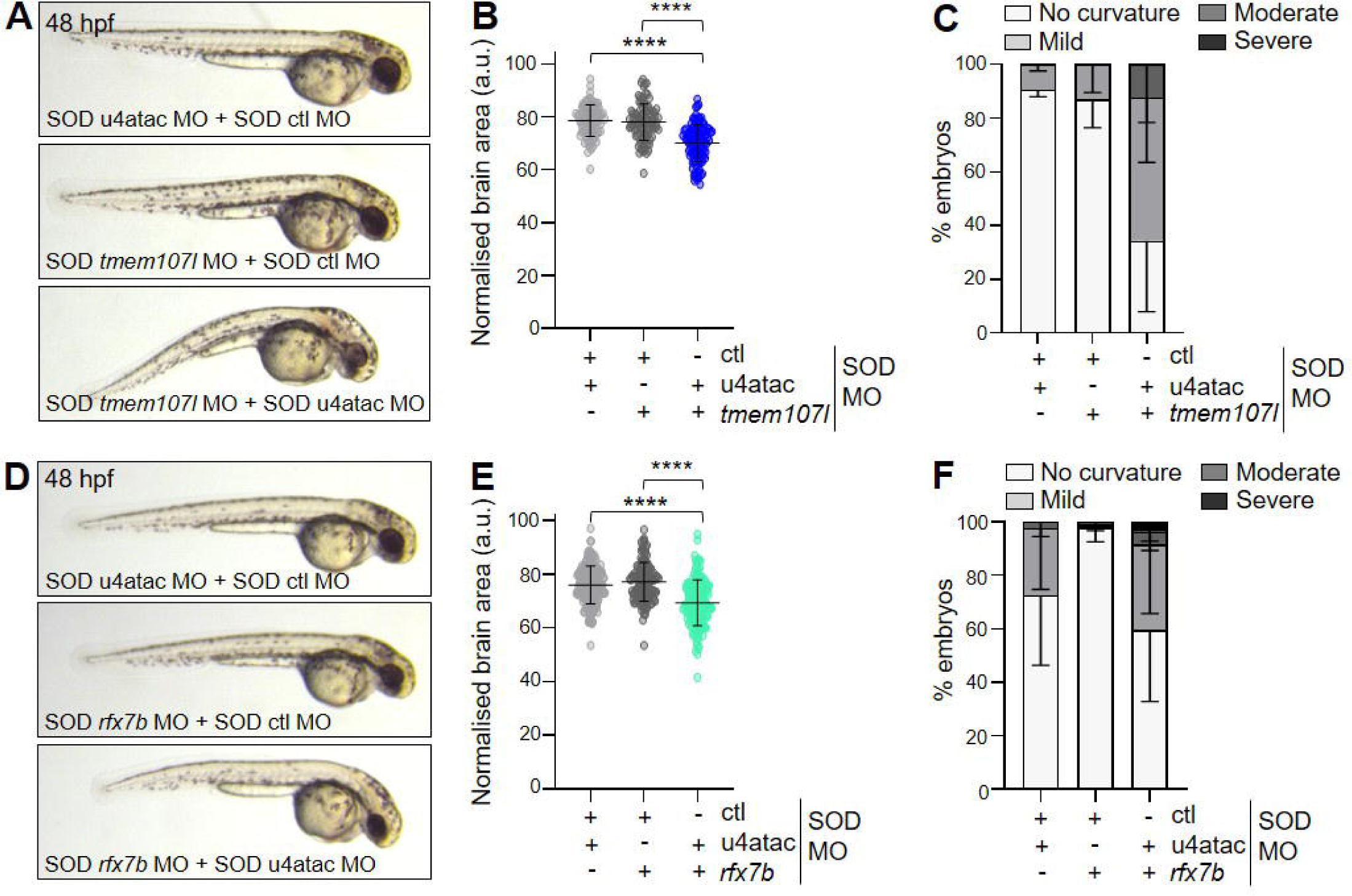
*tmem107l* and *rfx7b* genetically interact with *rnu4atac*. **(A, D)** Representative images of embryos injected with a combination of sub-optimal doses (SOD) of control MO, u4atac MO, or *tmem107l* (A) or *rfx7b* (D) U12 MO (as indicated on images) at 48 hpf. **(B, E)** Graphs showing the normalised head area to total body length at 48 hpf in the two epistasis conditions with either *tmem107l* (B) or *rfx7b* (E) SOD U12 MO. Graphs represent the mean ± SD of three independent experiments, with 30 to 60 embryos per experiment. ****P<0.0001, following one-way ANOVA test, with Dunnett’s comparison test. **(C, F)** Graphs showing the distribution of ventral body curvature phenotypes in the two epistasis conditions with either *tmem107l* (C) or *rfx7b* (F) SOD U12 MO at 48 hpf, categorized into four classes of severity (no curvature, mild, moderate, and severe) as shown in Supplementary Fig. 7F. Graphs show the mean ± SD from at least three independent experiments, each including a minimum of 30 embryos per batch.

We validated this experimental epistasis using a MO targeting *cep290,* a ciliary gene without U12-type intron that should not thus interact with *rnu4atac.* No genetic interaction was observed in animals injected with both *cep290* and u4atac MO SOD (Supp. Fig. 9A-C), whereas an optimal dose of *cep290* MO results in body curvature and mild microcephaly in zebrafish (Supp. Fig. 9D-F), as previously observed (Cardenas-Rodriguez et al., 2021). Furthermore, we checked whether an epistatic relationship could be observed with *tmem216*, encoding a ciliary transition zone protein that functionally interacts with *tmem107l* (Gogendeau et al., 2020; Lambacher et al., 2015). MO SOD for both genes led to the expected ciliary defects (body curvature, hydrocephaly), cardiac edema, and microcephaly (Supp. Fig. 9G-I), which were milder than the loss of function of *tmem216* (Supp. Fig. 9J-L). Hence, these control experiments validated the epistasis approach to tackle genetic interaction in zebrafish, and collectively, the data indicate that u4atac deficiency-associated ciliary traits and microcephaly are partially mediated by defective splicing of *tmem107l* and *rfx7b*.

#### TMEM107 *re-expression partially rescues MO-mediated deficiency of u4atac*

To validate the results obtained with the epistasis experiments, we investigated whether the observed phenotypes could be rescued by the injection of correctly spliced human *TMEM107* RNA.

First, we verified that human *TMEM107* RNA could replace that of zebrafish by performing rescue experiment in *tmem107l* U12 MO-injected embryos. Co-injection experiments resulted in the rescue of body curvature and microcephaly, while cardiac edema remained (Supp. Fig. 10A-B), suggesting that this latter trait may not be specific. We next performed co-injection of *TMEM107* RNA with u4atac MO. We validated in that context the impact of u4atac loss of function on endogenous *tmem107l* U12-type intron splicing by RT-qPCR analysis (Supp. Fig. 10C). While cardiac edema, otolith defects, and brain haemorrhages were observed in similar proportions to that detected in u4atac morphants (Figure 6A-B), the re-expression of *TMEM107* led to zebrafish with a much less severe axis curvature compared to the u4atac morphant condition (Figure 6A, C). The persistence of some of the phenotypes commonly associated with ciliary dysfunction indicates that other ciliary genes are involved as well in the ciliary phenotype of u4atac-deficient model, not surprisingly given the fact that there are 86 cilium-related MIGs. Nevertheless, we did observe that microcephaly was markedly reduced when *TMEM107* was re-expressed (Figure 6D), suggesting that the microcephaly observed in u4atac-deficient zebrafish embryos is, at least in part, attributable to mis-splicing of the U12-type intron of *tmem107l*, hence validating the observed results with the epistasis experiment.

**Figure 6.**
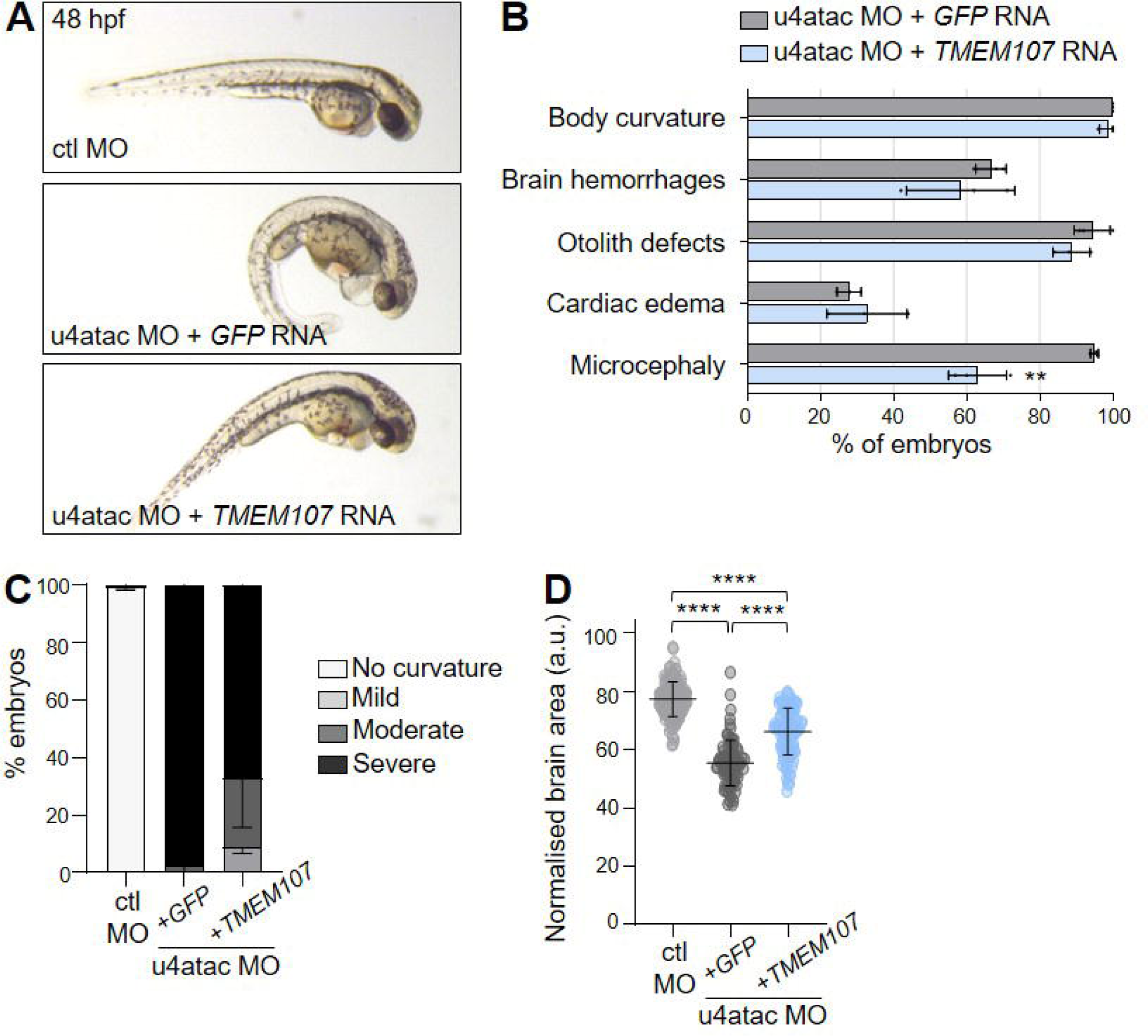
Human *TMEM107* mRNA partially restores microcephaly in u4atac-deficient zebrafish embryos. **(A)** Representative images of embryos at 48 hpf injected with control MO or u4atac MO in combination with either *GFP* or *TMEM107* RNA. **(B)** Diagrams representing the observed phenotypes in controls or *TMEM107*-rescued animals, as seen in A. Graph shows the mean ± SD of at least three independent experiments, with a minimum of 30 embryos per experiment. ns, non-significant, **P<0.01 following unpaired t-test. **(C)** Graph showing the distribution of ventral body curvature phenotypes in controls or *TMEM107*-rescued animals at 48 hpf, categorized into four classes of severity (no curvature, mild, moderate, and severe) as shown in Supplementary Fig. 6F. Graph shows the mean ± SD from at least three independent experiments, each including a minimum of 30 embryos per batch. **(D)** Graph showing the normalised head area to total body length at 48 hpf in controls and *TMEM107*-rescued animals. Graph represents the mean ± SD of three independent experiments, with 30 to 60 embryos per experiment. ****P<0.0001, following Kruskal-Wallis test, with Dunn’s comparison test.

## Discussion

In this study, we performed a genome wide transcriptomic analysis in u4atac-deficient zebrafish heads to unveil the molecular and cellular processes underlying the cerebral anomalies seen in *RNU4ATAC*-opathies. As expected, we detected massive U12-type intron retention with minimal effect on gene expression. Analysis of GO terms associated to DS U12-type introns or DE genes confirmed the central role of primary cilium. Two ciliary MIG were chosen for further exploration, and by combining epistasis and rescue approaches in zebrafish, we identified *TMEM107* as a functionally validated downstream effector of U4atac contributing to the neurodevelopmental defects associated with *RNU4ATAC*-related disorders, together with *RFX7* as an additional gene involved in microcephaly.

The present study is a follow-up of our previous work linking minor spliceosome with primary cilium function (Khatri et al., 2023). Since our publication, other studies confirmed this link. Bi-allelic mutations in two genes encoding protein components of the minor spliceosome, *SCNM1* and *ZRSR2*, both splicing factors associated to the U12 snRNP, have been associated to the Oro-Facial-Digital (OFD) ciliopathy (Hannes et al., 2024; Iturrate et al., 2022). Loss of function of these components leads to elongation of primary cilia in patients’ fibroblasts, and for *SCNM1*, to an alteration of Hh signalling, as we found in *RNU4ATAC* patients’ cells (Khatri et al., 2023). In addition, another U12 snRNP component, ZCRB1, has recently been shown to be a regulator of ciliogenesis and Wnt signaling in cellular and zebrafish models (Pirzada et al., 2026). Interestingly, *TMEM107* has been highlighted in these three studies as a cilium-related MIGs that exhibits either exon skipping (*SCNM1*), intron retention (*ZRSR2*) or decreased level of expression (*SCNM1*, *ZRCB1*), indicating that *TMEM107* gene is constantly altered in minor spliceosomopathies, which made it a good candidate to explore. It can also be mentioned that loss of function of *INTS13*, a subunit of the Integrator complex that is involved in the maturation of both major and minor snRNA (except U6 and U6atac), was associated to an OFD-like ciliopathy. Ciliogenesis was shown to be impaired in patients’ nasal cells, *INTS13*-deficient RPE1 cells, and in drosophila and xenopus models (Anderson et al., 2009; Jodoin et al., 2013; Mascibroda et al., 2020). Altogether, these data indicate that primary cilium biogenesis is a key cellular process regulated by minor splicing, either by a direct control of expression of ciliary U12-type intron-containing genes or indirect modulation of ciliary gene expression, as we observed in +16A and +51A zebrafish conditions. The latter situation could be attributed to the misregulation of ciliary gene expression programmes, mastered by specific transcription factors (TF) such as the *RFX* gene family. This TF class is known to bind a conserved motif, called X-box, that is largely present in the promoter regions of ciliary genes (Blacque et al., 2005; Swoboda et al., 2000). RFX5 and RFX7 are noticeable as they lack a dimerization domain, and are encoded by MIG. We observed that both gene transcripts present U12-type intron retention in our zebrafish dataset, and their loss of function may explain the misregulation of half of the down-regulated ciliary genes (40/80) that exhibit a X-box motif specific to their human counterparts.

Hence, by picking both *TMEM107* and *RFX7* as candidate MIG to explain part of the ciliopathy-related phenotypes seen in u4atac zebrafish model, we were confounded to observe that the zebrafish orthologues mostly contributed to the microcephaly phenotype. Albeit microcephaly is not a cardinal feature of ciliopathies, it is not rare to observe reduction of brain size in patients with Bardet-Biedl, Joubert or OFD syndromes (reviewed in Thomsen et al., 2025). This outcome can be explained either by the pleiotropic functions of microcephaly-related proteins, or by the tight dependence of cilium assembly/disassembly with cell cycle. In the first case, there are few examples of key regulators of mitotic spindle organization that also participate to cilium assembly or length (KIF11, Zalenski et al., 2020; RTTN, Vandervore et al., 2019), while loss of function of CPAP, WDR62 or RRP7 results in delayed cilium disassembly, causing cell cycle entry retardation and overall decrease proliferation of neuronal progenitors (Farooq et al., 2020; Gabriel et al., 2016; Zhang et al., 2019). Hence, for TMEM107, we can hypothesize that it contributes to brain size through its core function at the transition zone of primary cilia, known to guarantee the correct regulation of the developmental Shh signalling pathway, important for neuronal progenitor expansion (Wang et al., 2016). For RFX7, *CCND2* and *TUBA1A* were shown to be putative direct gene targets (Schwab et al., 2023), both of which being involved in human neurodevelopmental disorders (Bahi-Buisson et al., 2008; Mirzaa et al., 2014).

Beyond the role of primary cilium in *RNU4ATAC*-opathy physiopathology, it is noteworthy that our transcriptomic dataset points to other altered cellular processes. Notably, it is striking that loss of function of u4atac (u4atac MO, +16A, +51A) results in dysregulation of genes involved in immune response. This observation is in line with the immunodeficiency seen in patients (Cuinat et al., 2026; Lovric et al., 2026), and with the recent study in a drosophila model of u4atac loss of function. Authors showed that u4atac depletion leads to splicing defects in transcripts involved in innate immunity and haematopoiesis, with a special focus on JAK/STAT signalling (Shikara et al., 2026). In addition to the known roles of certain MIG in the regulation of innate immune response (such as PARP1 and ZDHHC18, Supp. Dataset 1), we can cite, here again, the role of RFX7, which has been implicated in the natural killer cell maintenance (Castro et al., 2018) or in the response to cytomegalovirus infection by controlling SOCS3 expression (Wang et al., 2023). To support this hypothesis, we observed hematopoietic defects in *rfx7b* morphants, as revealed by O-dianisidine staining (Supp. Fig. 8B, C).

Finally, in this study, we chose to compare two common *RNU4ATAC* variants that are each associated to either a severe (TALS, n.51G>A, 51 reported cases) or milder (RFMN, n.16G>A, 11 reported cases) form of *RNU4ATAC*-opathies. The DE genes or DS introns we detected in our zebrafish RNAseq datasets were basically the same for these two variants, thus preventing us to draw any conclusions or hypotheses regarding distinct physiopathological mechanisms that would explain the severity difference between n.51G>A and n.16G>A. Considering more carefully DSI, it was counterintuitive to observe that U12-type intron retentions affected more transcripts and were of higher level in +16A than in +51A condition, in line with our former work (Khatri et al., 2023). More significant was the difference between variants and the u4atac MO condition. We interpreted this difference by the hypomorphic nature of the variants. It has to be mentioned that we also observed a remarkable difference in the splicing of U2-type introns, which seemed better spliced in presence of *RNU4ATAC* variants compared to controls. This effect of U4atac deficiency on U2-type intron splicing was also observed in RFMN patients’ mononuclear blood cells (Cologne et al., 2019) and in TALS twins’ lymphoblastoid cells (Almentina Ramos Shidi et al., 2022). We hypothesised that the U2-type introns concerned by this phenomenon could be those usually physiologically retained through alternative splicing, these IRs playing a key role during cell development, cell differentiation, and in response to cellular stress (Boutz et al., 2015; Braunschweig et al., 2014; Grabski et al., 2021). This finding supports an unforeseen potential interplay between the minor and major spliceosomes, that could perhaps give a “raison-d’être” to the maintenance of minor splicing in nearly all species (Moyer et al., 2020). Further experiments are needed to better characterize the role of U4atac in the regulation of physiological U2-type retained introns, and to determine whether alteration of this novel role of U4atac participate to the physiopathological mechanisms underlying *RNU4ATAC*-opathies.

## Acknowledgments

We thank the AZR aquatic core facility (Charlotte Perret, Annie Désenfant, Olivier Lohez), and GenCyTi platform for their assistance. We also thank Alizée Lemasson for her technical inputs and all the GENDEV team members for stimulating discussions. This work was supported by CNRS, Inserm and Université Lyon 1 through recurrent funding, the Agence Nationale de la Recherche (ANR-18CE12-0007-01 and ANR-22-CE12-0007-01), the Fondation Maladies Rares for transcriptomic analyses (FONDATION-GenOmics_202003011), and Fondation pour la Recherche sur le Cerveau-Rotary for the confocal microscope. C.J. was supported by the ANR (ANR-22-CE12-0007-01) and the Fondation pour la Recherche Médicale (FDT202504020193).

## Competing interests

The authors report no competing interests.

## Notes

### Competing Interest Statement

The authors have declared no competing interest.

