## Supplemental information for "u4atac regulates cilium biogenesis through splicing of the minor intron of *tmem107l* and *rfx7b* in zebrafish developing brain"

This PDF file includes:

Supplementary Figures 1 to 10 and legends

Supplementary Table 1

SI Reference

**Supplementary figures and legends**

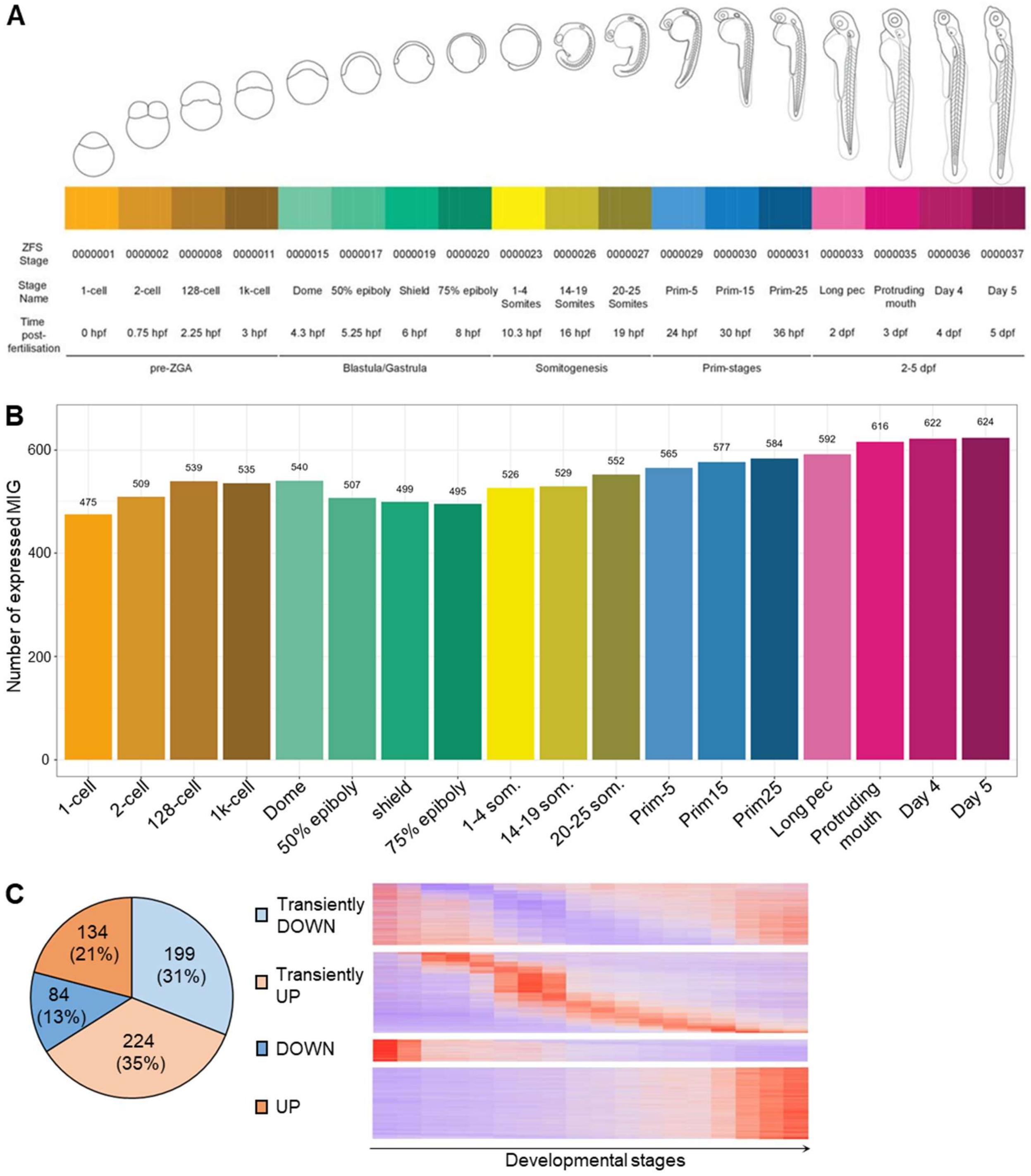


**Supplementary Figure 1. Minor intron-containing genes are largely expressed during early development in zebrafish. (A)** Schema issued from White et al. 2017 depicting all the 18 developmental stages analysed by RNAseq. Five replicates for each time point were done. **(B)** Diagram indicating the number of MIG expressed (mean TPM ≥ 1) for each developmental stage, out of the 677 we identified in the GRCz11 genome assembly. **(C)** Diagram showing the distribution of the 641 expressed MIG (left) depending of their expression pattern: transiently down, transiently up, progressively down or progressively up throughout the developmental stages, with low expression shown in purple and high expression in red (right).

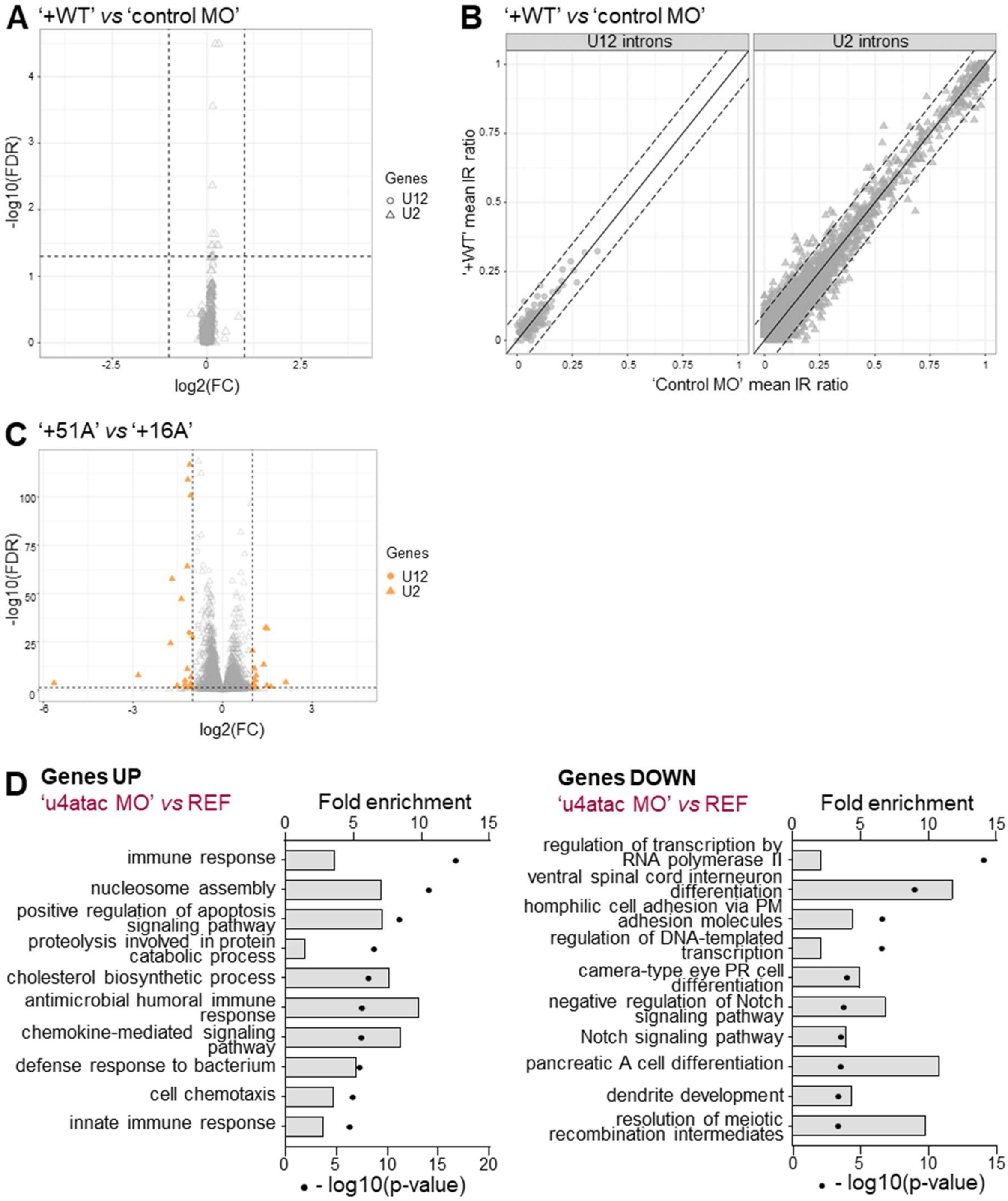


**Supplementary Figure 2. Human U4atac snRNA fully restores u4atac loss of function in zebrafish. (A)** Volcano plot of U2 and U12 gene expression in +WT condition compared to control MO condition. U12 genes are represented as dots and U2 genes as triangles. Dashed lines delineate the significance boundaries (FDR = 0.05 and |log2(FC)| = 1), with upregulated genes represented in red, downregulated in blue, and those not meeting the threshold ($P < 0.05$, FDR < 0.1, > ± 2 FC) in grey (NS). **(B)** Distribution of U12-type and U2-type intron retention levels in +WT condition relative to control MO condition. The solid diagonal line indicates equal retention levels between the two conditions (y = x). Dashed lines delineate a ±10% difference threshold relative to the equality line. Grey dots/triangles correspond to introns with non-significant differences. **(C)** Volcano plot of U2 and U12 gene expression in +51A condition compared to +16A condition. U12 genes are represented as dots and U2 genes as triangles. Dashed lines delineate the significance levels (FDR = 0.05 and |log2(FC)| = 1). Grey dots/triangles correspond to genes with non-significant differences. **(D)** GO term enrichment analysis of up- and down-regulated genes in the u4atac MO condition compared with the REF condition, ranked according to the adjusted p-value (black dots). Bars represent fold enrichment for each GO term.

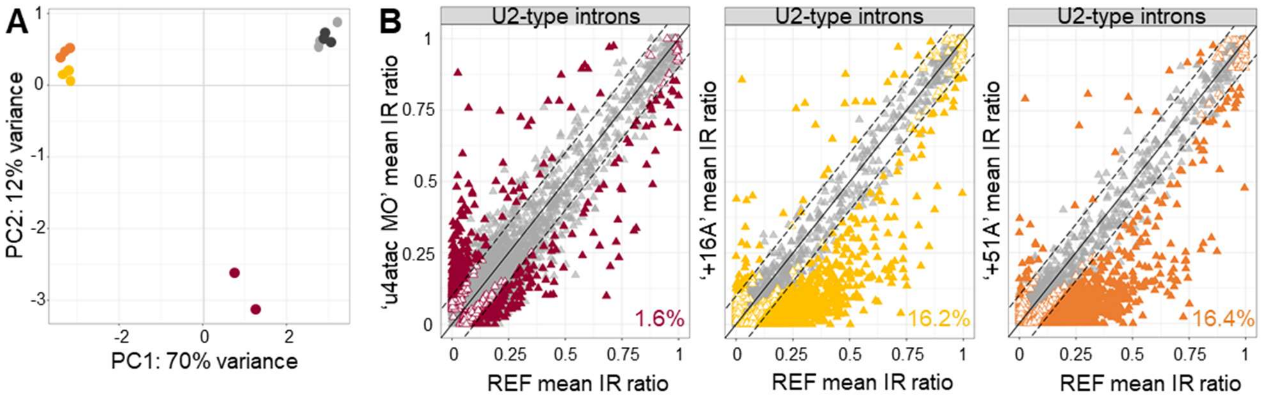


**Supplementary Figure 3. A significant proportion of U2-type introns are better spliced in U4atac-16A or -51A snRNA expressing embryos. (A)** PCA of U2-type intron retention across the five experimental conditions, based on the 500 most variable introns. Color legend is shown in Figure 1B. **(B)** Distribution of U2-type intron retention levels in each condition relative to the reference (REF). All 11 808 detected U2-type introns are shown. The solid diagonal line indicates equal retention levels between the two conditions (y = x). Dashed lines indicate a ±10% difference threshold relative to the equality line. Colored triangles indicate statistically significant differences (FDR < 0.05), with plain triangles being introns with ΔIR ratio > 0.1 and empty triangles with ΔIR ratio < 0.1, while grey triangles correspond to introns with non-significant differences (FDR > 0.05). Percentages indicate numbers of U2-type introns with significant |ΔIR ratio| > 0.1 over all significant U2-type introns.

**A**

| Specie | Gene | Genomic position | RefSeq or Ensembl ID | Protein length | Amino acid identity (%) in functional domains |
|---|---|---|---|---|---|
| Homo sapiens | *TMEM107* | 17: 8,172,457 - 8,176,380 | NM_183065.4 | 140 aa | |
| | *RFX7* | 15: 56,087,280 - 56,245,082 | NM_022841.7 | 1 460 aa | |
| Danio rerio | *tmem107* | 14: 247,456 - 251,089 | ENSDARG00000059150.1 | 135 aa | 57,4 |
| | *tmem107-like* | 23: 28,482,780 - 28,487,153 | ENSDARG00000088398.1 | 140 aa | **65,4** |
| | *rfx7a* | 7: 34,506,937 - 34,546,833 | ENSDARG00000077237.2 | 1 509 aa | 86,8 |
| | *rfx7b* | 25: 387,996 - 405,745 | ENSDARG00000103691.1 | 1 551 aa | **91,2** |

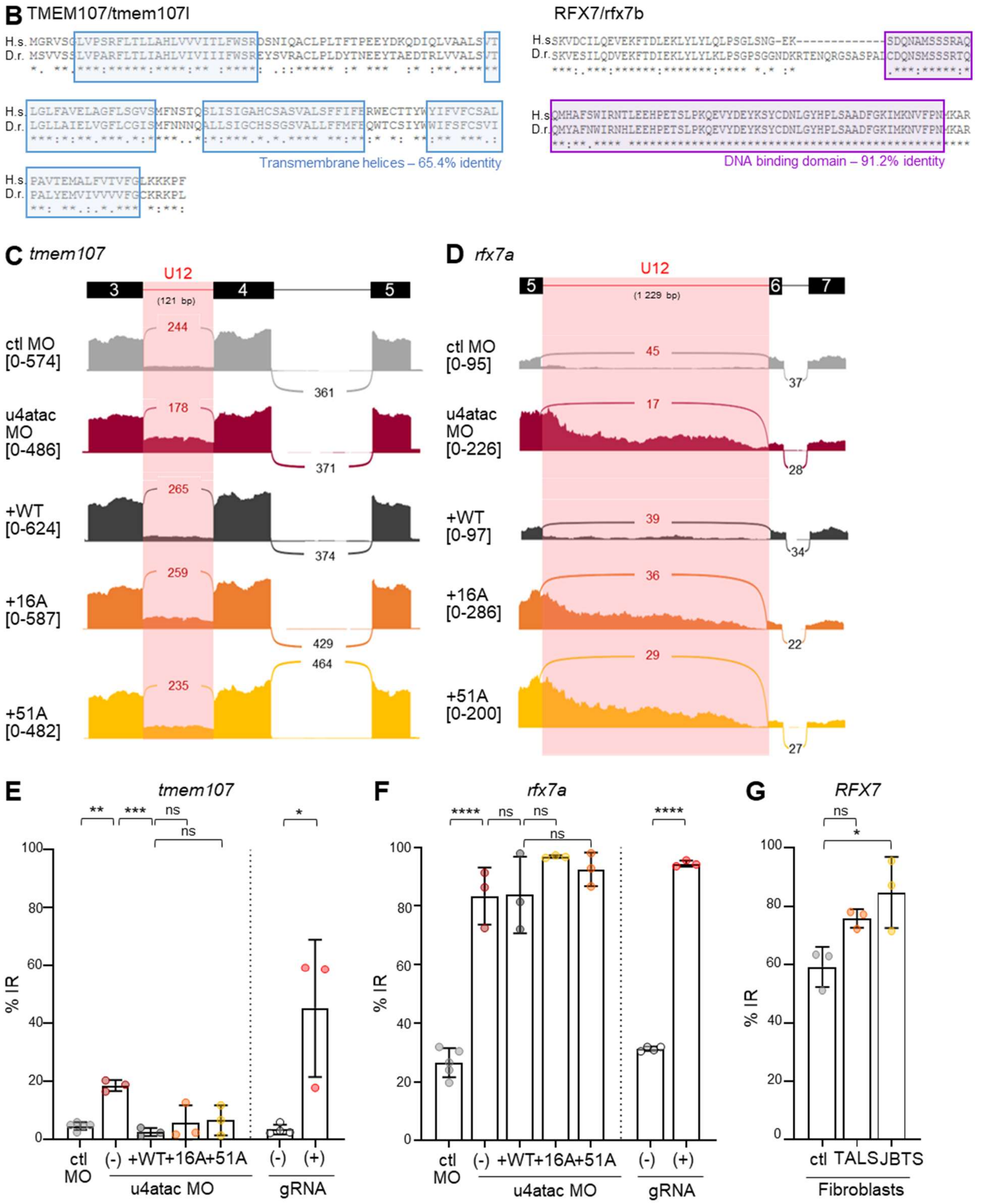

**Supplementary Figure 4. *TMEM107* and *RFX7* orthologues in zebrafish and analysis of U12-type intron retention of *tmem107* and *rfx7a* paralogues. (A)** Table listing the human and zebrafish orthologues of *TMEM107* and *RFX7* genes with their genomic position, RefSeq or Ensembl ID, and length of the encoded protein. For each tmem107 and rfx7 zebrafish paralogue, the percentage of amino acid identity with the human protein for the functional domains is indicated. **(B)** Protein sequence alignment of the human (H.s.) and zebrafish (D.r.) orthologues of TMEM107 (left, full protein shown) and RFX7 (right, portion of the protein comprising the only known functional domain). Known protein domains are highlighted by colored boxes. Stars indicate identical residues. **(C-D)** Sashimi plots of *tmem107* (C) and *rfx7a* (D) across the different experimental conditions, focused on the region encompassing the U12-type intron (highlighted in pink). On lines, the number of reads split across splice junctions; in brackets, the range of coverage of each base of the depicted region. **(E-G)** RT-qPCR quantification of *tmem107* (E) and *rfx7a* (F) U12-type intron retention in 48 hpf embryos across the different conditions. In G, quantification of human *RFX7* U12-type intron retention in patient's fibroblasts with Taybi-Linder (TALS) or Joubert-like (JBTS) syndromes described in Khatri et al. (2023). Graphs show the mean ± SD of at least three independent experiments. ns, non-significant, *P<0.05, **P<0.005, ***P<0.001, ****P<0.0001 using one-way ANOVA test with Dunnett's multiple comparisons (MO conditions, human samples), or unpaired t-test (gRNA conditions).

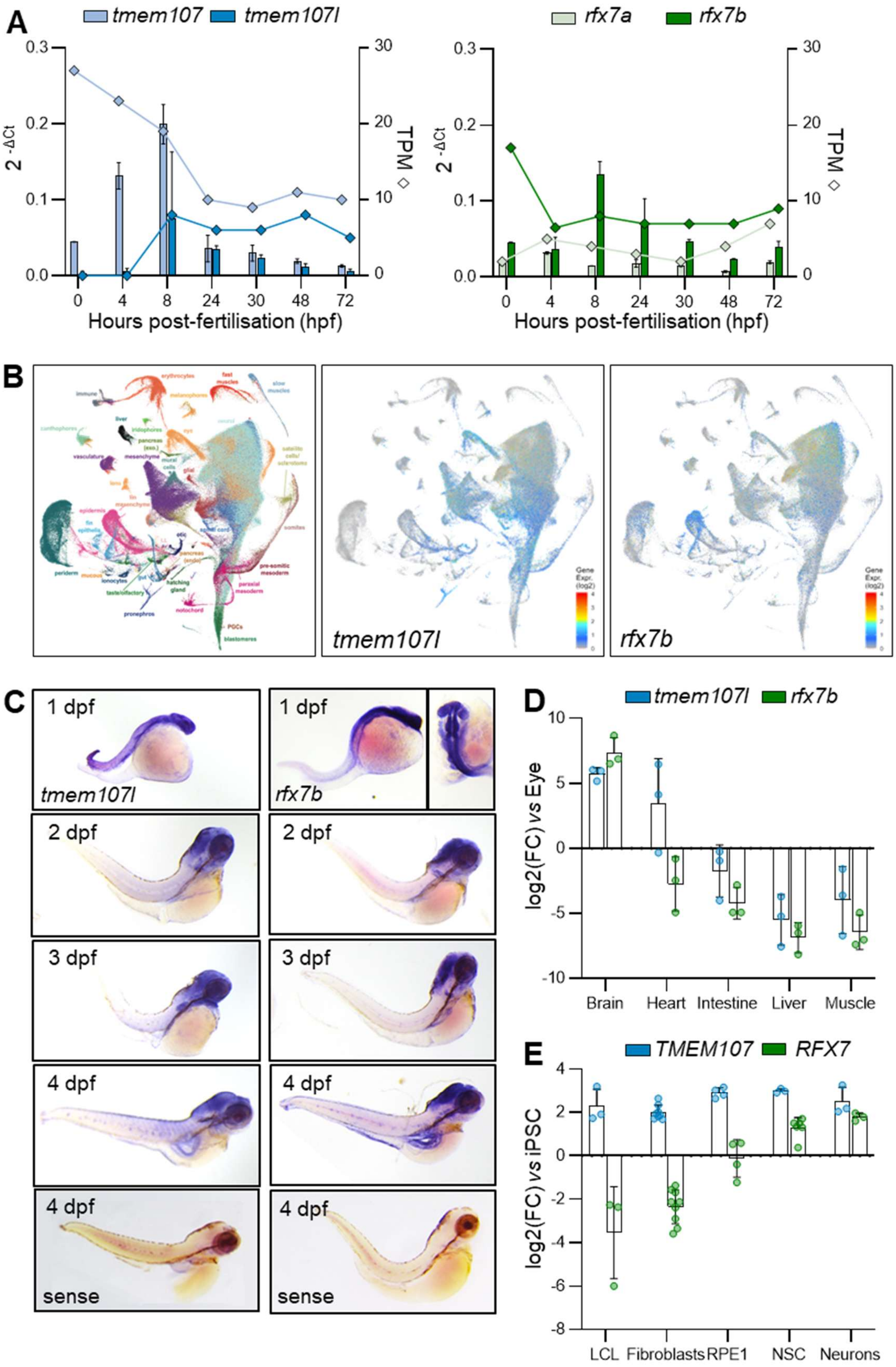


**Supplementary Figure 5. *tmem107l* and *rfx7b* are predominantly expressed in the developing zebrafish brain. (A)** Graphs showing *tmem107*/*tmem107-like* (left) and *rfx7a/b* (right) gene expression during zebrafish early embryo development (0 to 72 hpf). On the left Y-axis, values of

RT-qPCR data calculated with $2^{-\Delta Ct}$ method and represented by bars (mean ± SD), and on the right Y-axis, TPM values issued from RNAseq dataset of White et al. 2017 and represented by diamonds. **(B)** Expression of *tmem107l* and r*fx7b* across zebrafish cell types during early development (1 to 5 dpf) using DanioCell resource (https://daniocell.nichd.nih.gov/, Farrell et al. 2018; Sur et al. 2023). **(C)** Whole-mount *in situ* hybridization analysis of *tmem107l* and *rfx7b* expression from 1 to 4 days post-fertilisation (dpf). Sense probes were used as negative controls. **(D)** RT-qPCR analysis of *tmem107l* and *rfx7b* expression in adult zebrafish tissues (4 months post-fertilization). Graph shows the mean ± SD of three biological replicates; values were normalized to the expression level of the eye. **(E)** RT-qPCR analysis of *RFX7* and *TMEM107* expression in human cell types. Graph shows the mean ± SD of three biological replicates; values were normalized to the expression level in induced pluripotent stem cells (iPSCs).

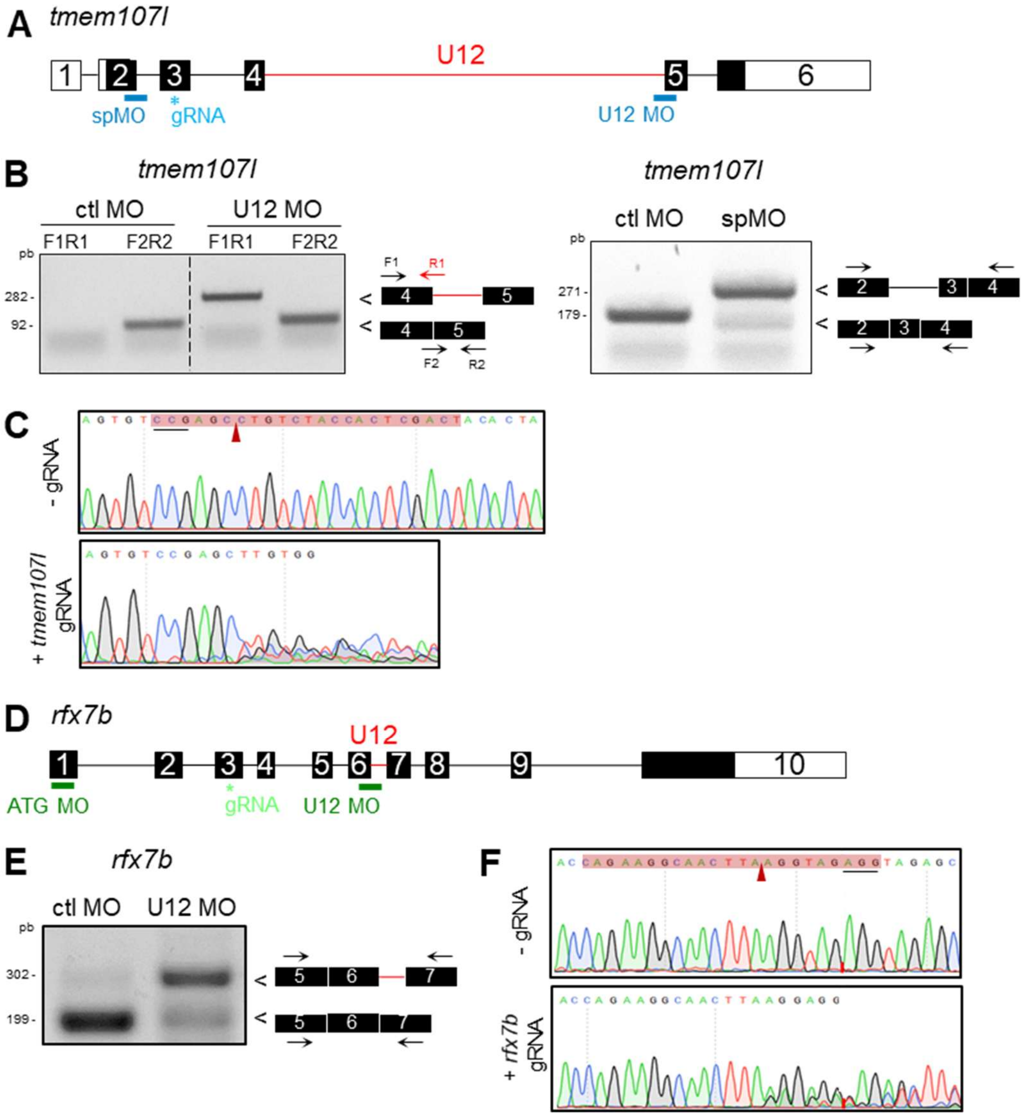


**Supplementary Figure 6. Validation of MO and CRISPR/Cas9 RNP efficiency to alter *tmem107l* and *rfx7b* expression. (A, D)** Schematic representation of *tmem107l* (A) and *rfx7b* (D) gene structures. Exons are represented as boxes and introns as connecting lines. The U12-type intron is highlighted in red, and coding exons are shown in black. The binding sites of the splice (sp), ATG and U12-type intron (U12) morpholino oligonucleotide (MO), and of CRISPR/Cas9 guide RNA (gRNA), are indicated. **(B, E)** RT-PCR validation of *tmem107l* (B) and rfx7b (E) mis-splicing following U12 (B-left panel, E) or splice (B-right) MO injection compared to the control condition. All MO induce intron retention. Primer positions are indicated. **(C, F)** Representative chromatograms depicting *tmem107l* (C) and *rfx7b* (F) gRNA target region in control (- gRNA) and crispant (+ gRNA) embryos. The sequence of gRNA is highlighted in red, with the PAM sequence underlined and the red arrow showing the double stand break position.

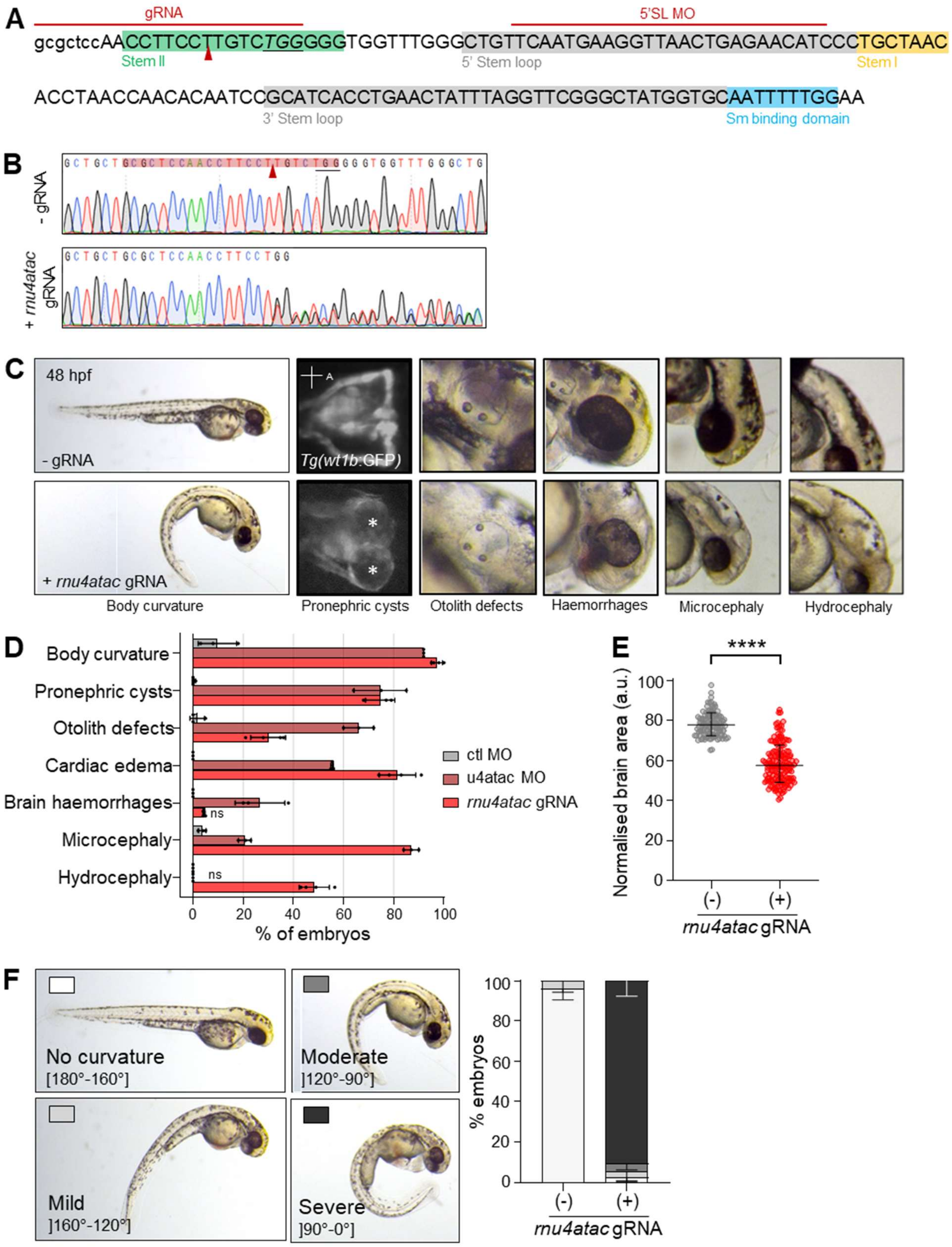


**Supplementary Figure 7. CRISPR/Cas9-mediated loss of function of *rnu4atac* induces severe developmental defects in F0 mosaic zebrafish embryos. (A)** Schematic representation of the zebrafish *rnu4atac* sequence showing the positions of 5'SL MO and CRISPR/Cas9 RNP (gRNA) target sites. Known *rnu4atac* functional regions are indicated by colored boxes. The red triangle indicates the predicted CRISPR/Cas9 cleavage site. **(B)** Representative chromatograms depicting the *rnu4atac* gRNA target region in control (- gRNA) and crispant (+ gRNA) embryos. The sequence of gRNA is highlighted in red, with the PAM sequence underlined and the red arrow showing the double stand break position. **(C)** Representative images of observed phenotypes at 48 hpf in control condition and crispant embryos. From left to right: body curvature, pronephric cysts (shown by asterisks in dorsal view, anterior to the right), otolith defects, haemorrhages (and microphthalmia), microcephaly, and hydrocephaly. **(D)** Percentage of embryos displaying each of the observed phenotypes in crispants compared with control and *u4atac* 5'SL morphants (Khatri et al. 2023). Graph shows the mean ± SD of at least three independent experiments, with a minimum of 30 embryos per experiment. ****P<0.0001, following two-way ANOVA with Tukey's multiple comparison test unless stated otherwise (ns, non significant). **(E)** Graph showing the normalised head area to total body length in *rnu4atac* crispants at 48 hpf. Graph represents the mean ± SD of three independent experiments, with 30 to 60 embryos per experiment. ****P<0.0001, following Mann-Whitney test. **(F)** Left panel, representative images illustrating ventral body curvature classes depending on the angle drawn between the head, the yolk extension and tip of the tail of zebrafish embryos at 48 hpf. Right panel, graph showing the distribution of ventral body curvature phenotypes as seen in *rnu4atac* crispants in left panel. Graph shows the mean ± SD of at least three independent batches of a minimum of 30 embryos.

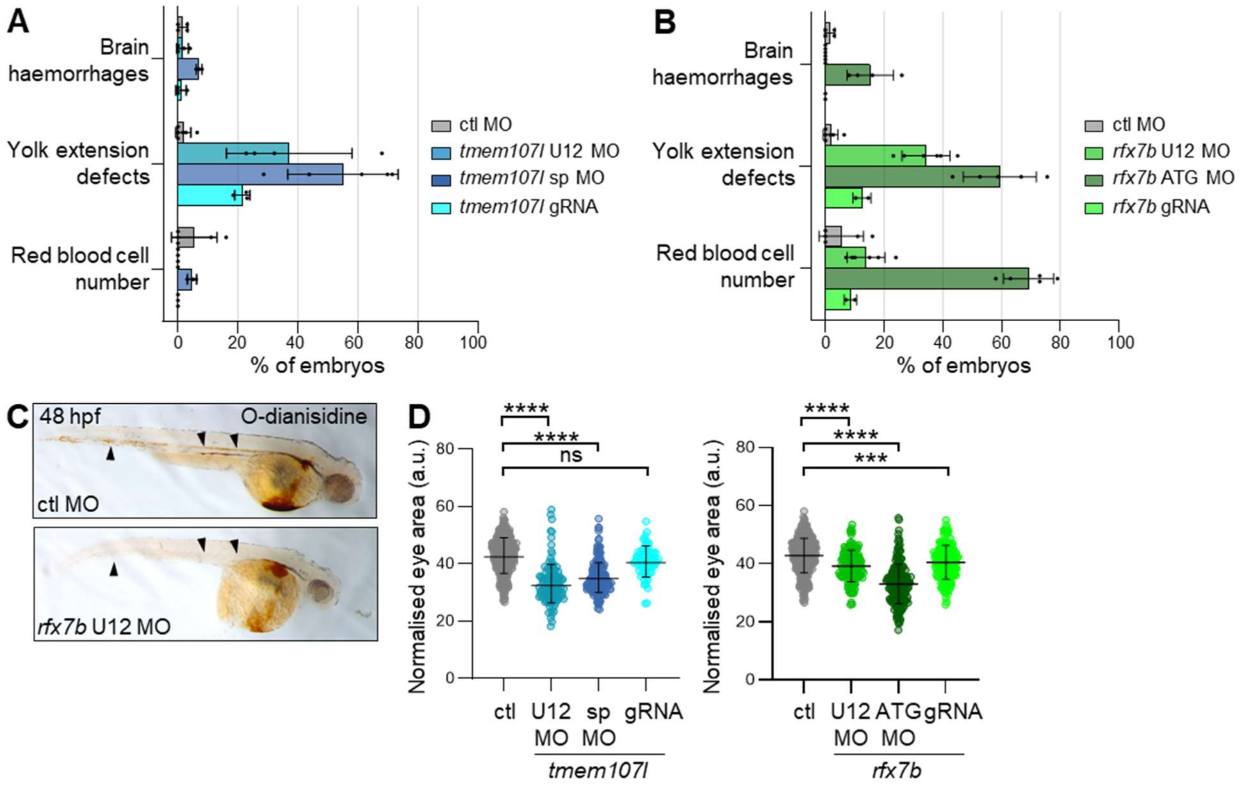


**Supplementary Figure 8. Additional phenotypes observed in *tmem107l* and *rfx7b* deficient animals. (A-B)** Percentage of embryos displaying each of the additional phenotypes observed in *tmem107l* (A) and *rfx7b* (B) deficient conditions (splice (sp) MO or ATG MO, U12 MO, or gRNA) compared with control morphants. Graphs show the mean ± SD of at least three independent experiments, with a minimum of 30 embryos per experiment. **(C)** Representative images of O-dianisidine staining on control or *rfx7b* MO-injected embryos at 48 hpf. Black arrowheads point to blood cells in the control animal, absent in *rfx7b* morphant embryo. **(D)** Graphs showing the normalised eye area to total body length in *tmem107l* (left) or *rfx7b* (right) deficient embryos at 48 hpf. Graphs represent the mean ± SD of three independent experiments, with 30 to 60 embryos per experiment. ***P<0.001, ****P<0.0001, following Kruskal-Wallis test with Dunn's multiple comparison test.

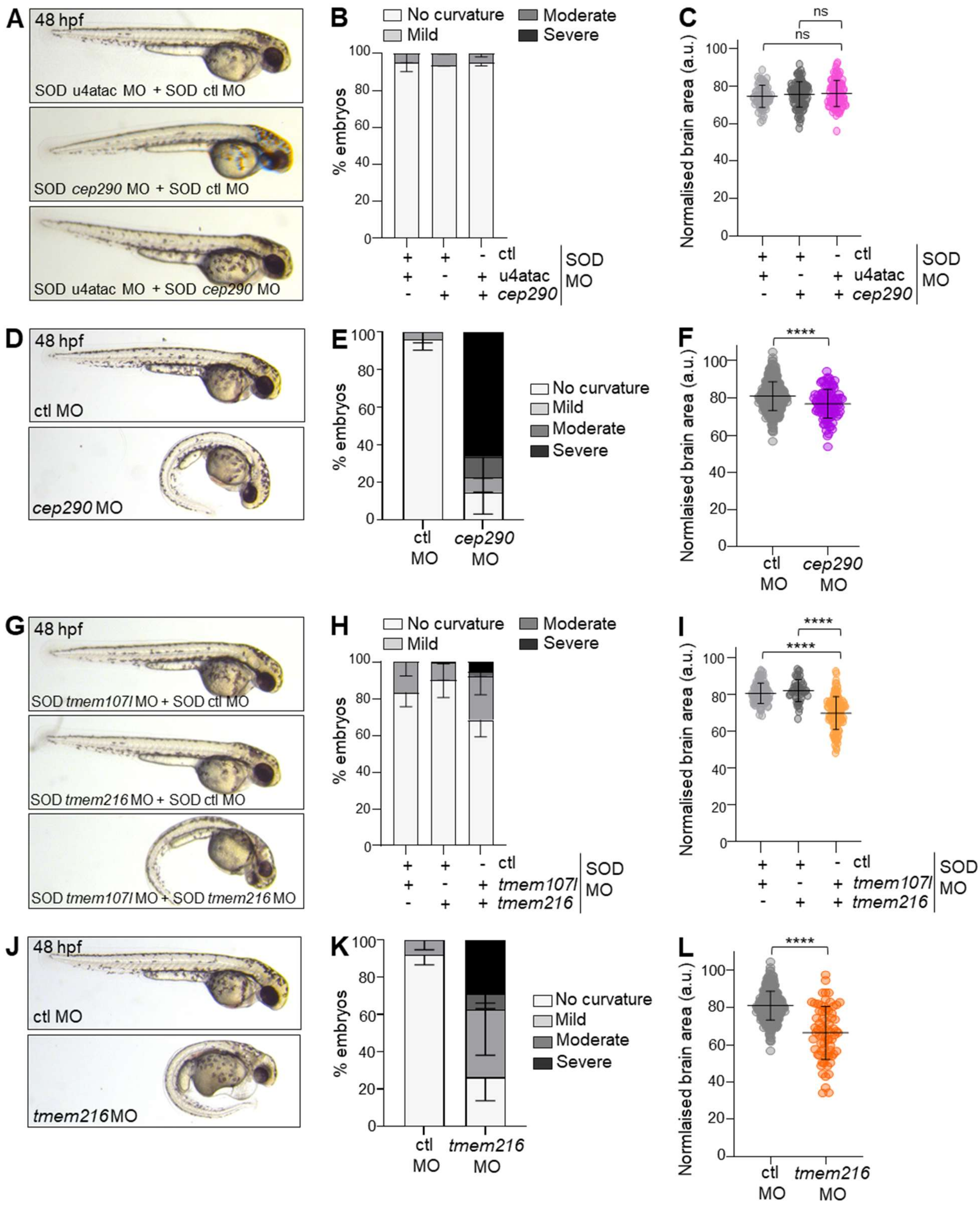


**Supplementary Figure 9. Validation of epistasis approach using *cep290* and *tmem216* as controls**. **(A, G)** Representative images of embryos injected with combination of sub-optimal doses (SOD) of control MO, u4atac MO, and *cep290* MO (A) or of control MO, *tmem107l*, and *tmem216* MO (G) (as indicated on images) at 48 hpf. **(B, H)** Graphs showing the distribution of ventral body curvature phenotypes in the two epistasis conditions with either u4atac and *cep290* (B) or *tmem107l*

and *tmem216* (H) SOD MO at 48 hpf, categorized into four classes of severity (no curvature, mild, moderate, and severe) as shown in Supplementary Fig. 7F. Graphs show the mean ± SD from at least three independent experiments, each including a minimum of 30 embryos per batch. **(C, I)** Graphs showing the normalised head area to total body length at 48 hpf in the two epistasis conditions with either u4atac and *cep290* (C) or *tmem107l* and *tmem216* (I) SOD MO. Graphs represent the mean ± SD of three independent experiments, with 30 to 60 embryos per experiment. ns, non-significant, ****P<0.0001, following one-way ANOVA test, with Dunnett's comparison test (C) or Kruskal-Wallis test with Dunn's comparison test (I). **(D, J)** Representative images of embryos injected with control MO or *cep290* MO (D) or *tmem216* MO (J) at 48 hpf. **(E, K)** Graphs showing the distribution of ventral body curvature phenotypes in control or *cep290* (E) or *tmem216* (K) MO-injected embryos at 48 hpf, categorized into four classes of severity (no curvature, mild, moderate, and severe) as shown in Supplementary Fig. 7F. Graphs show the mean ± SD from at least three independent experiments, each including a minimum of 30 embryos per batch. **(F, L)** Graphs showing the normalised head area to total body length at 48 hpf in control or *cep290* (F) or *tmem216* (L) MO-injected embryos at 48 hpf. Graphs represent the mean ± SD of three independent experiments, with 30 to 60 embryos per experiment. ****P<0.0001, following Mann-Whitney test.

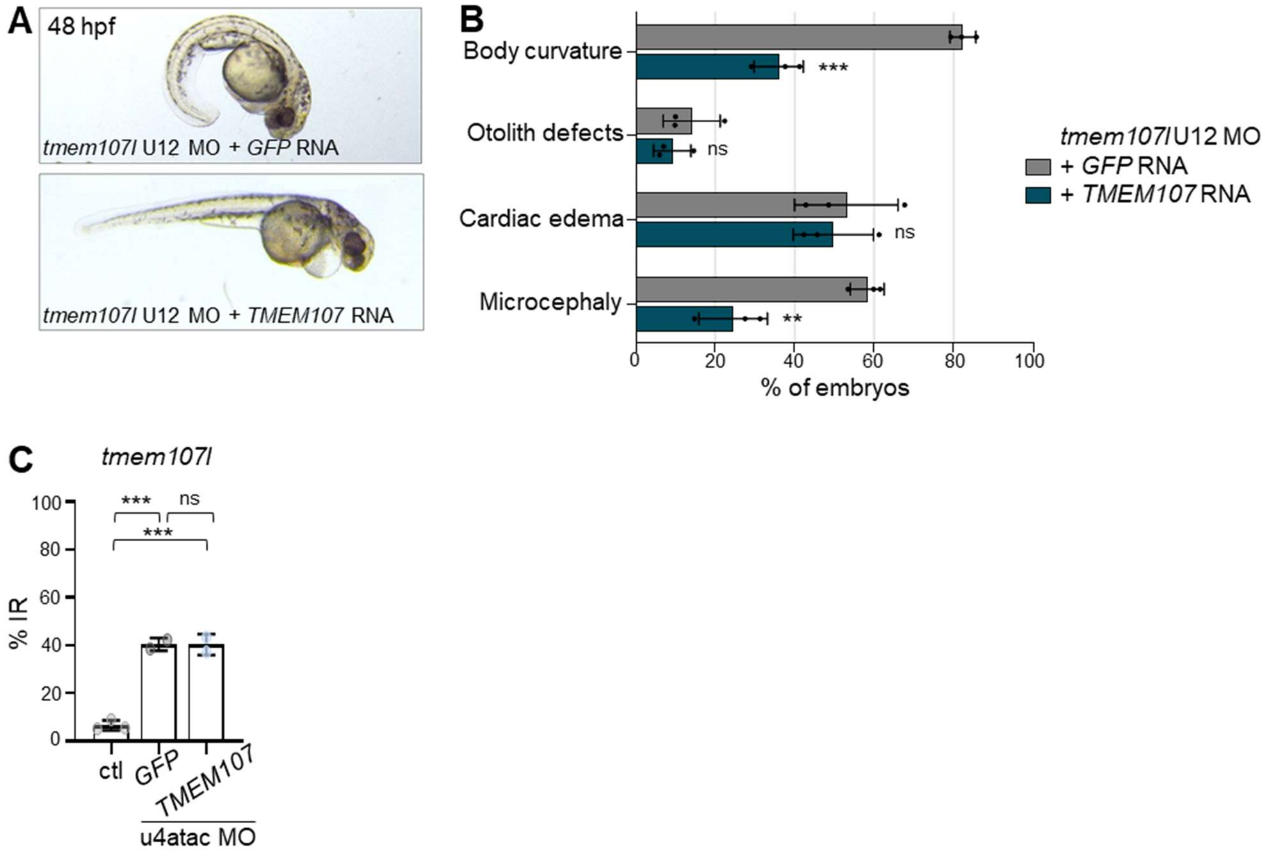


**Supplementary Figure 10. Validation of rescue approach of *TMEM107* RNA in *tmem107l* MO-injected embryos. (A)** Representative images of embryos at 48 hpf injected with *tmem107l* U12 MO in combination with either GFP or human *TMEM107* RNA. **(B)** Percentage of embryos displaying each of the phenotypes observed in *TMEM107*-rescued *(+ TMEM107* RNA*)* animals compared to control (+ *GFP* RNA). Graph shows the mean ± SD of at least three independent experiments, with a minimum of 30 embryos per experiment. ns, non-significant, **P<0.01, ***P<0.001, following unpaired t-tests. **(C)** Graph of RT-qPCR analysis of *tmem107l* U12-type intron retention showing the mean ± SD of at least two independent experiments. ns, non-significant, ***P<0.001 using one-way ANOVA test with Dunnett's multiple comparisons.

**Supplementary Table 1. Primer, morpholino oligonucleotide and crRNA sequences**

**RT-PCR primer sequences**

| Gene | Target sites | Primer F | Primer R |
|---|---|---|---|
| *rfx7b* | Exon5 – Exon7 | CAGTGGTGGCAATGACAAAAG | CTCCACACTGTCAAACTGGC |
| | Exon6 – Exon7 | TTATTGCGACAATCTGGGTTA | CCCATCCCCTGATTTGTGTA |
| *tmem107l* | Exon2 – Exon4 | ATGTCGGTGGTCAGCAGT | GTGTGTCCTCTGCTGTGTACT |
| | Exon4 – Intron4 | ACCAAGCTCTTCTGTGTATCCT | GCTGTGACATCCAATAGCTGA |

**RT-qPCR primer sequences**

| Gene | Isoform(s) | Primer F | Primer R |
|---|---|---|---|
| *rfx7b* | unspliced U12-type intron | TCCTTTACAATGCATCACATACTG | TGTAAGTCCAGGTTAGGGAGC |
| | spliced U12-type intron | GGCATGAGGGGGAAATCAAAATAC | CCCATCCCCTGATTTGTGTA |
| | all | TTATTGCGACAATCTGGGTTA | AACGCCGTGCCTTCATATTC |
| *tmem107l* | unspliced U12-type intron | TTATTTAACCGTCACTTTTCAGCTA | AAATGCTGCAGGTCCATTGC |
| | spliced U12-type intron | CAACAACCAAGCTCTTCTGTC | GCAGGTCCATTGCTCAAACAT |
| | all | TCATTTTCTGGTCTCGGGAGT | GTGTGTCCTCTGCTGTGTACT |
| *rfx7a* | unspliced U12-type intron | GCGAGGCAAATCCAAATATCCT | ACACAGAGTTCAGGGACAGG |
| | spliced | CTCTCAGTGCTGCGGACTTT | TGGCATCTGTACAAAGGTCTTC |
| | all | AAATCGGGTGATGGGTGTGA | TTCAACGCTGTCAAACTGCC |
| *tmem107* | unspliced U12-type intron | GTTTCTGAACGTCCGTCAGCTC | AGCAGGGCCACTGGTGAAAA |
| | spliced U12-type intron | GTAATCAGGCCCTGCTGTCTCT | GATGATCCAGTACGTCCAGCAG |
| | all | AACAGAAGACACACGGCTGA | ACTCCAGACAGAAACCCAGC |
| *gapdh* | all | GTTGGTATTAACGGATTCGGT | CACTTAATATTGGCTGGGCT |
| *RFX7* | unspliced U12-type intron | GGTTAAAGCCCTTGTGACACT | GCCAGCCACCAGTAACAAAC |
| | spliced U12-type intron | TTCCAAACATGAAGGCACGT | GTCCACTGTAGCAATATTTAG |
| | all | TGTGACCAGAGGACCAAATGT | TGCTGCTGGTGAAAGTTAAGT |
| *TMEM107* | unspliced U12-type intron | CACCACTGGCCTTTTCTGAC | ATGAAGAAGGACAGGGCCAC |
| | spliced U12-type intron | GCACCCAGAGCCTCATCTC | CACTCCCAACGCTCGAATATG |
| | all | CGCGCTCTCTGTCACCCT | GGGTGCTGTTGAACATGGAG |
| *RPS17* | all | CATTATCCCCAGCAAAAAGC | AGGCTGAGACCTCAGGAACA |

**Primer sequences for ISH probes**

| Gene | ISH probe | Primer F[a] | Primer R[a] |
|---|---|---|---|
| *rfx7b* | antisense | TCGAAAAGCTCTACCTCTACCTTAA | TAATACGACTCACTATAGGGTTTAGGTAAAATCTGGGCATATCGG |
| | sense | TAATACGACTCACTATAGGGTCGAAAAGCTCTACCTCTACCTTAA | TTTAGGTAAAATCTGGGCATATCGG |
| *tmem107l* | antisense | AAGCTTGATAAACTTATACGCATAGTAGCC | TAATACGACTCACTATAGGGCAAAATGCCTTTAATCATCACAAGAGGTTC |
| | sense | TAATACGACTCACTATAGGGAAGCTTGATAAACTTATACGCATAGTAGCC | CAAAATGCCTTTAATCATCACAAGAGGTTC |

**MO sequences**

| Transcript | Target site | Sequence |
|---|---|---|
| 5-mismatch control | | 5'- GATcTTgTCAcTTAACCTTgATTcA -3' |
| u4atac | 5' stem-loop | 5'- GATGTTCTCAGTTAACCTTCATTGA -3' |
| *rfx7b* | ATG | 5'- TGTTGTTGATCATCGGCCAT -3' |
| | Exon6-Intron6 (U12-type) junction | 5'- TGACCACAGAAAAGGATATTTTGAT -3' |
| *tmem107l* | Exon2-Intron2 (U2-type) junction | 5'- AGAACACTTCTTACCCGAGACCAG -3' |
| | Intron4 (U12-type)-Exon5 junction | 5'- AGAACACTTCTTACCCGAGACCAG -3' |
| *tmem216* | ATG | 5'- TTCCGTGGGCAGCCATGTTAGATTG -3' |
| *cep290* | ATG | 5'- TTGATGTGTACCAGTTGTGCTGATG -3' |

**crRNA sequences**

| Gene | Sequence[b] |
|---|---|
| *rnu4atac* | 3'- GCGCTCCAACCTTCCTTGTC *TGG* -5' |
| *rfx7b* | 3'- CAGAAGGCAACTTAAGGTAG *AGG* -5' |
| *tmem107l* | 3'- AGTCGAGTGGTAGACAGGCT *CGG*-5' |

[a]T7 promoter underlined
[b]PAM sequence in italic
